# PhenoDEL as a Novel Screening Strategy Based on Intracellular Protein Degradation Activity

**DOI:** 10.1101/2025.11.26.690606

**Authors:** Yuichi Onda, Yurika Ochi, Toshihiro Araki, Miho Kageoka, Shuzo Takeda, Kazunori Yamada, Takehiko Ueda, Ken Ohno, Minoru Tanaka, Daiki Sakai, Miki Hasegawa, Yoshihito Tanaka

## Abstract

Targeted protein degradation (TPD), including proteolysis targeting chimeras (PROTACs) and molecular glue degraders (MGDs), is a promising therapeutic approach. However, systematic discovery of such small molecules remains a major challenge. Here, we present PhenoDEL, a novel phenotypic DNA-encoded library (DEL) screening platform that integrates one-bead one-compound DEL (OBOC-DEL) with the Beacon® optofluidic system for high-throughput, single-cell analysis. By co-culturing individual OBOC-DEL beads and engineered reporter cells in nanoliter-scale chambers, PhenoDEL enables direct observation of compound-induced protein degradation at single-cell resolution. We demonstrate this approach by identifying compounds that induce degradation of FKBP12^F36V^-EGFP fusion proteins in PC-3 cells. The workflow allows precise linkage between compound identity and cellular phenotype via DNA barcoding and next-generation sequencing. PhenoDEL overcomes limitations of conventional screening methods, offering high sensitivity, spatial control, and scalability. This platform holds significant potential for mechanism-driven drug discovery, including identification of novel PROTACs and MGDs.

**Graphical abstract:** 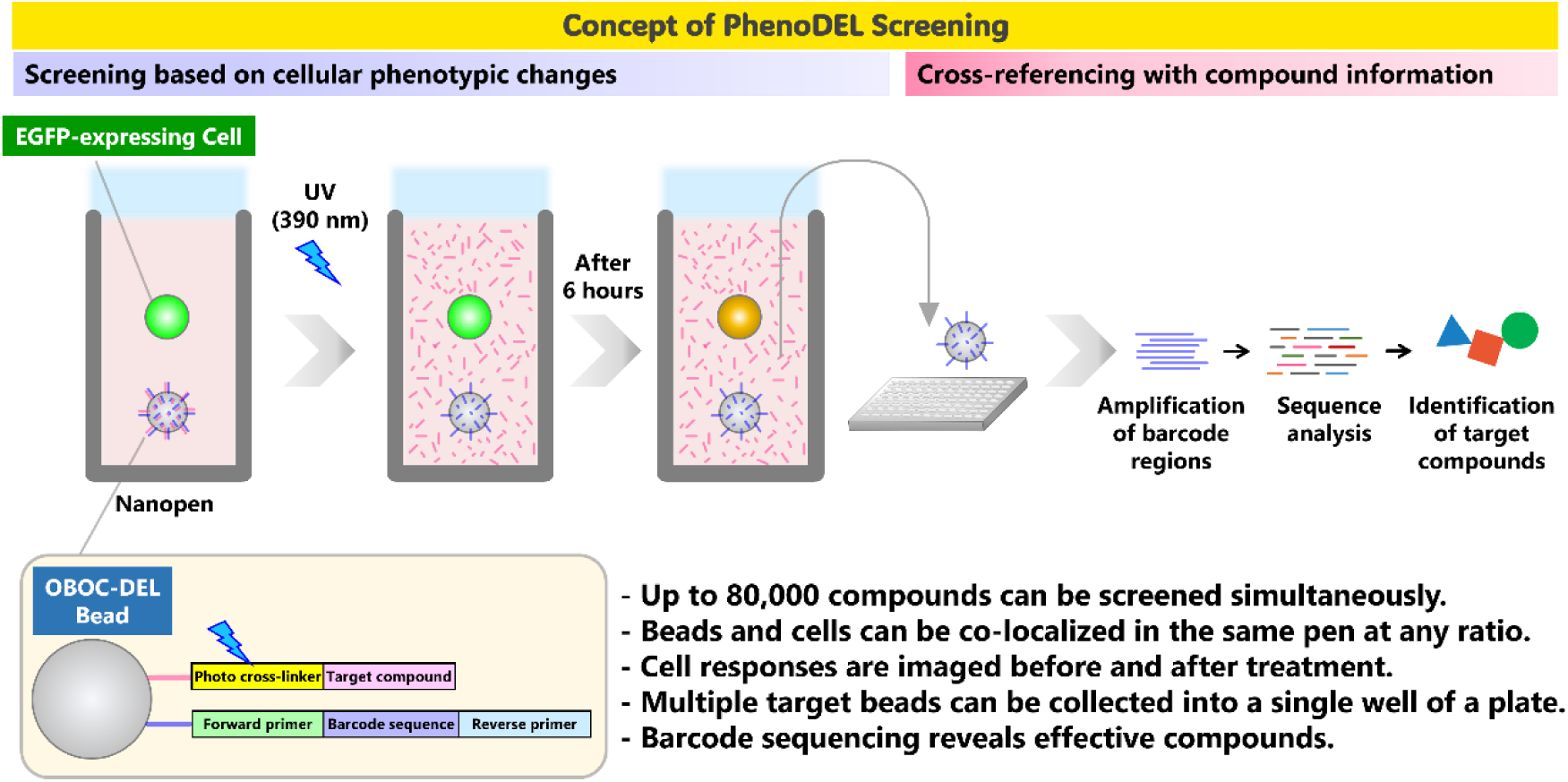

## Introduction

Targeted protein degradation (TPD) is emerging as a promising approach in new drug discovery and development, utilizing technologies like proteolysis targeting chimeras (PROTACs) and molecular glues degraders (MGDs) ^1^ . The systematic discovery of small molecules that induce proximity between proteins enables the targeted modulation and degradation of previously undruggable proteins, thereby expanding the scope of drug discovery to include a wide range of biological processes and molecular targets through event-driven pharmacology ^2^ . PROTACs are heterobifunctional molecules that recruit E3 ligases (E3L) to degrade protein of interest (POI), while MGDs facilitate protein-protein interactions between POI and E3L to induce degradation. Although most MGDs have historically been discovered serendipitously through phenotypic or viability-based screens, recent advances in mechanistic understanding and assay technologies are enabling a shift toward more systematic and rational approaches for their identification^3^. Accordingly, rational design and high-throughput screening (HTS) are becoming essential to improve the efficiency and specificity of MGD discovery, ultimately enhancing their therapeutic potential.

DNA encoded library (DEL) technology can evaluate a vast number of compounds (millions to billions) at once, enabling the discovery of ligands for undruggable targets that are difficult to target using conventional drug discovery methods such as HTS^4^ . DEL has been used not only for the discovery of binders, but also for the optimization of compounds, especially PROTACs that induce protein degradation, as well as for the discovery of MGDs^5^. Recently, solid-phase DEL which is called one-bead one-compound DEL (OBOC-DEL) has been developed to further expand DEL screening from affinity screening to activity screening^6^. (Figure 1A). Paegel and co-workers have reported an integrated droplet-based microfluidic circuit that directly screens OBOC-DEL (67,100 compounds) against phosphodiesterase autotaxin (ATX) for enzyme inhibitors^6c^. Paegel and Paulick have reported in vitro transcription–translation (IVTT) activity assay with GFP reporters fused to target genes, screening for inhibitors using OBOC-DEL (5,348 compounds) in microfluidic droplets and the hits inhibited PCSK9-GFP IVTT and reduced PCSK9 levels in HepG2 cells^7^. Scientists at Plexium and Amgen have described Picowell RNA-seq screening to identify The Von Hippel-Lindau (VHL) MGDs from a diverse VHL-focused OBOC-DEL (4,327 compounds), discovering dGEM3, which targets GEMIN3 for degradation, highlighting the potential of VHL for novel target identification^8^.

**Figure 1.**
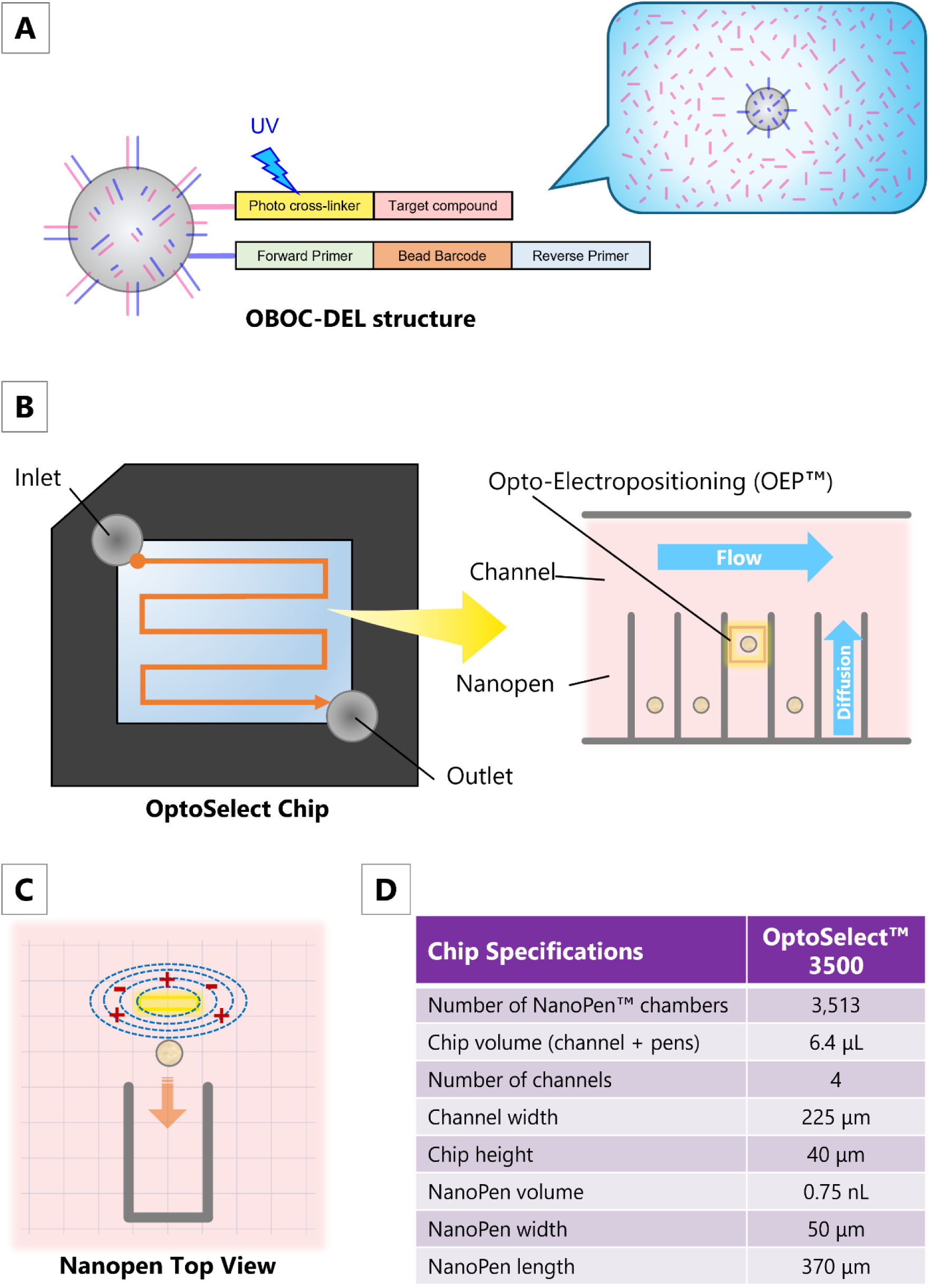
(A) One-bead one-compound DEL (OBOC-DEL) structure. The linker is cleaved by irradiation with near-ultraviolet light (390 nm) and the compound is released into the solution. (B) The OptoSelect™ chip used in the Beacon® Optofluidic System (#110-08004 /Bruker Corporation) has a structure called a NanoPen™ chamber, which has a volume approximately 100,000 times smaller than a microwell, and a flow path called a channel, which are arranged on a semiconductor substrate. (C) Particles in the liquid loaded into the flow channel are “pushed” into the nanopenby a light-induced electric field using Opto-Electropositioning (OEP™). Liquid exchange within the nanopenis achieved by diffusion. OEP optical technology non-destructively irradiates the OptoSelect Chip substrate with visible light to create a localized electric field gradient. Cells and particles in liquid are usually repelled by this electric field gradient. The Beacon system can move cells and particles within the chip by controlling the light pattern and electric field gradient. (D) The OptoSelect™ 3500 Chip (#750-00012/Bruker Corporation) used in our experiment has approximately 3,500 nanopenswith a volume of 0.75 nL. Up to four chips can be run simultaneously, theoretically allowing screening of more than 10,000 samples at a time.

The Beacon® optofluidic system (currently distributed by Bruker Cellular Analysis) integrates optics, fluidics, and imaging to enable high-throughput single-cell manipulation and analysis^9^. Its OptoSelect™ chip contains 3,500 NanoPen™ chambers (∼0.75 nL each), ∼100,000 times smaller than microwells, allowing screening of over 10,000 samples (Figure 1B, 1D). Cells are positioned using Opto-Electropositioning (OEP™), a light-induced electric field for precise, non-invasive control (Figure 1C).

Initially developed for cell line development, Beacon achieves >99% single-cell origin verification, outperforming limiting dilution and fluorescence-activated cell sorting^10^. In antibody discovery, it enables direct screening of antibody-secreting cells via bead-based fluorescence assays, identifying high-affinity clones without hybridoma generation^11^. It also supports multiplexed functional assays for CAR-T and TCR-T cells, linking phenotype to genotype at single-cell resolution^9b,12^. While Beacon excels in screening large biomolecules like antibodies, applying it to small molecules requires additional engineering. Specifically, compounds must be individually delivered and retained in nanopen chambers to ensure localized exposure. To address this, we developed PhenoDEL—a phenotypic DNA-encoded library screening method that leverages Beacon’s single-cell resolution and compartmentalized architecture to identify small molecule degraders.

In this study, we introduce a novel screening concept termed PhenoDEL which combines OBOC-DEL technology with single-cell phenotypic analysis on the Beacon platform. As a proof-of-concept, we applied PhenoDEL to screen for compounds that induce degradation of FKBP12^F36V^-EGFP fusion proteins in engineered PC-3 cells. This approach enables direct observation and evaluation of cellular responses before and after compound release at the single-cell level, which is not achievable with conventional technologies. By comparing cellular phenotypes before and after the assay and excluding cells with low initial activity, PhenoDEL reduces the likelihood of false positives and allows for the accurate identification of active degraders.

## Results

### Concept of single bead screening in Beacon

To enable phenotypic screening at single-cell resolution, we developed a workflow that integrates OBOC-DEL beads with FKBP12^F36V^-EGFP reporter cells using the Beacon® Optofluidic System. In this workflow, each NanoPen™ chamber on the OptoSelect™ chip was loaded with a single OBOC-DEL bead and a single PC-3 cell genetically engineered to express FKBP12^F36V^-EGFP fusion proteins (Figure 2). After loading, residual liquid in the channel was removed by CO₂ flow, and compounds were then released from the beads by near-ultraviolet (UV-A) irradiation through a DAPI filter (excitation wavelength: 390/40 nm). The released compound diffused into the nanopen and interacted with the co-cultured cell. Changes in EGFP fluorescence were monitored to evaluate compound-induced phenotypic effects. Beads associated with cells exhibiting EGFP fluorescence loss were collected and subjected to next-generation sequencing (NGS) to identify the compound via its DNA barcode. This single-bead, single-cell co-culture strategy enabled a direct link between compound identity and cellular phenotype, providing a powerful platform for activity-based screening of DNA-encoded libraries.

**Figure 2.**
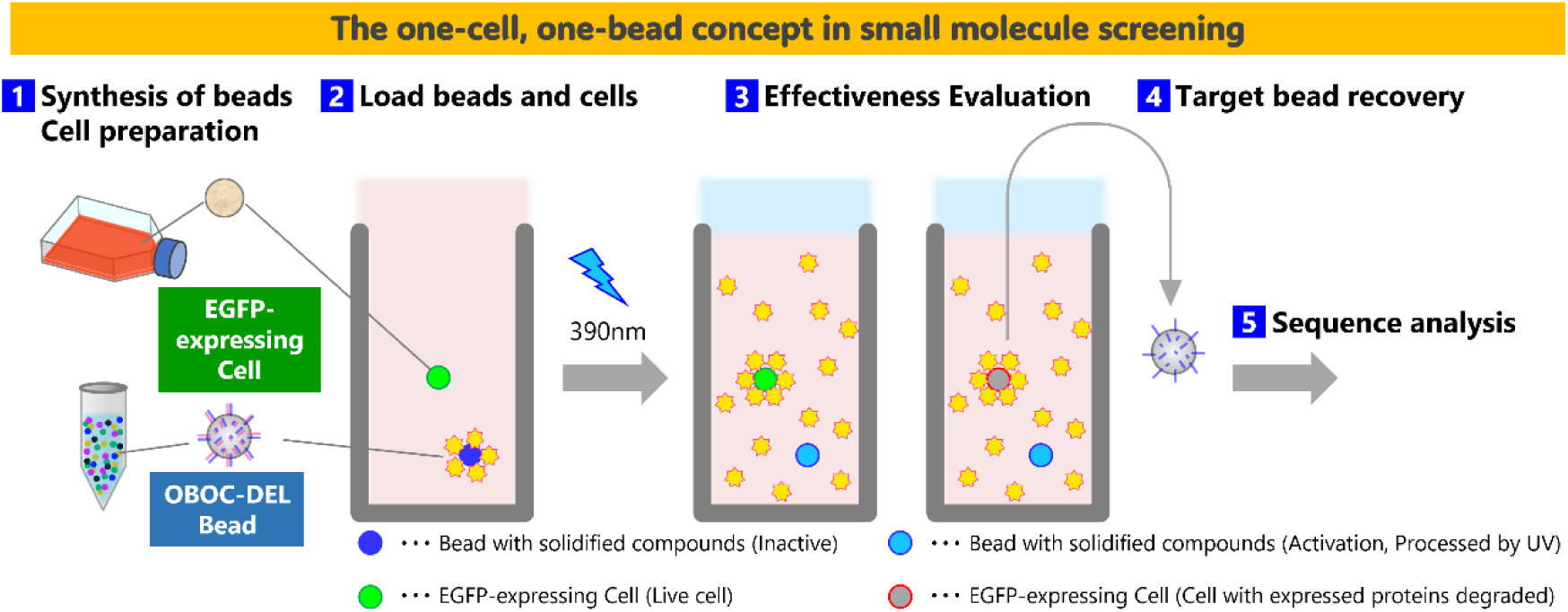
Concept of single bead screening on the Beacon platform: (1) OBOC-DEL beads synthesized for this experiment were suspended at a concentration of 1.5 × 10⁶ particles/mL in PBST. PC-3 cells engineered to express FKBP12^F36V^-EGFP fusion proteins were cultured for at least two passages to ensure acclimation and adjusted to 2.0 × 10⁵ cells/mL immediately prior to loading into the Beacon system. (2) A single bead was loaded into each nanopen, followed by the introduction of a single FKBP12^F36V^-EGFP-expressing PC-3 cell into the same chamber. After successful penning of both the bead and the cell, residual liquid in the channels was removed by CO₂ flow for a defined period. (3) Near-ultraviolet (UV-A) light was irradiated through a DAPI filter (excitation wavelength: 390/40 nm) for 200 ms to cleave the compound from the bead. The released compound diffused into the nanopen and interacted with the co-cultured cell. Time-lapse imaging was performed to monitor changes in EGFP fluorescence, which served as a readout for compound-induced phenotypic effects. (4) Beads from nanopen in which EGFP fluorescence had disappeared were unloaded and collected into a 96-well plate. (5) The DNA barcode attached to each bead was amplified by PCR and analyzed via NGS to identify the compound. Barcode sequences were mapped to specific compound structures.

### Optimized protocol for compound release from OBOC beads using the Beacon system

To facilitate high-resolution phenotypic screening on the Beacon® Optofluidic System, we developed and optimized a protocol for compound release from fluorescein-conjugated OBOC beads (Figure 3A). These beads were synthesized using a photolinker strategy and served as a model system to evaluate the efficiency of linker cleavage and compound diffusion within nanoliter-scale NanoPen™ chambers.

**Figure 3.**
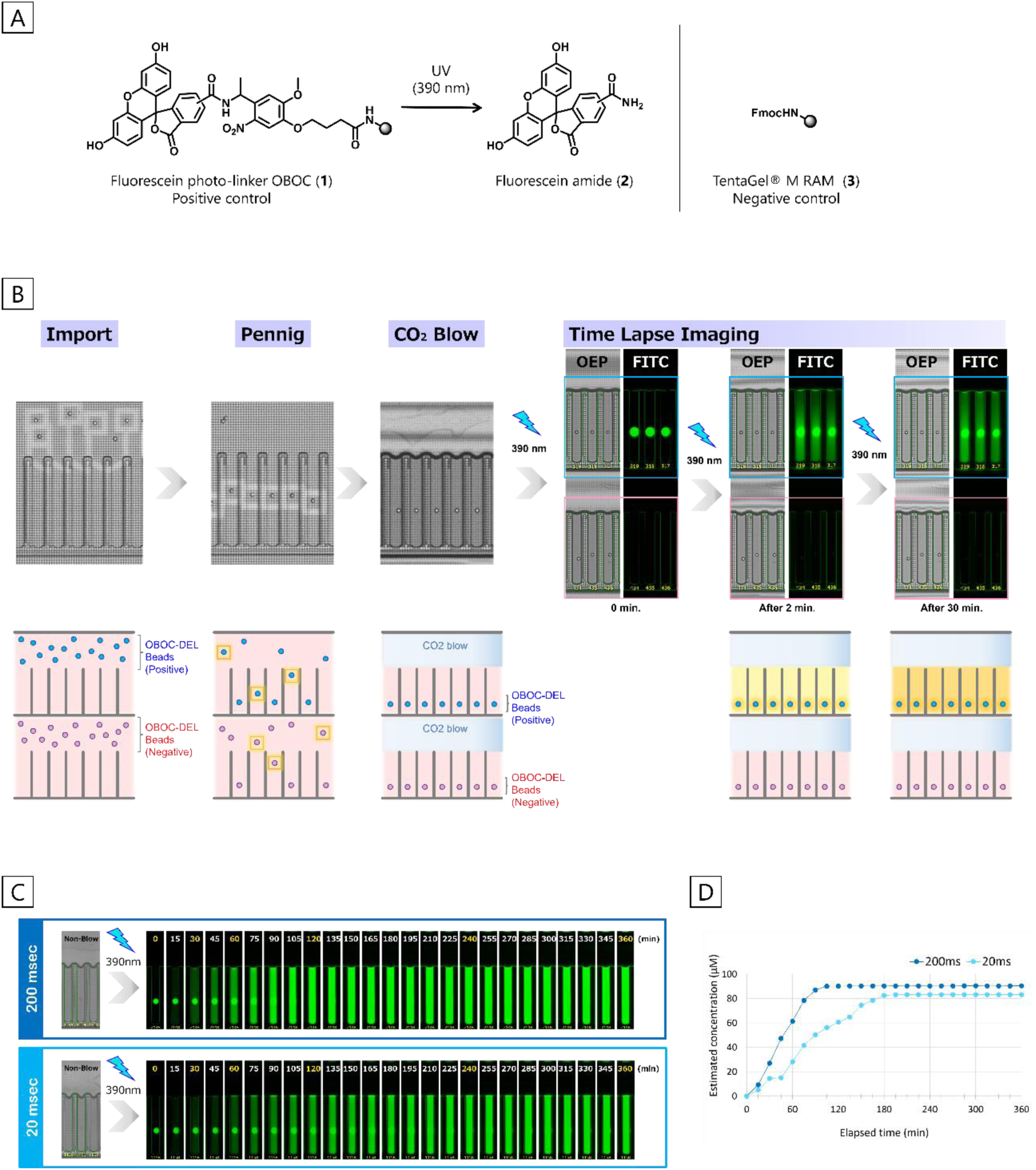
Procedure of the linker cleavage experiment using Beacon with OBOC beads. (A) Schematic of linker cleavage induced by 390 nm light through a DAPI filter. In the validation experiments, the beads were conjugated with a photo-cleavable linker labeled with fluorescein. (B) OBOC beads adjusted to 1.5×10⁶ particles/mL were loaded into the OptoSelect™ chip and individually penned into nanopens using Opto-ElectroPositioning (OEP). Residual liquid in the channel was removed by manual CO₂ flow. Linker cleavage was induced by irradiation with 390 nm light through a DAPI filter, applied for 10 seconds per cycle. FITC fluorescence from released compounds was monitored at defined intervals using time-lapse imaging, with each imaging session preceded by irradiation. In positive beads (top row), FITC fluorescence released from the linker was observed at 2 minutes. In contrast, negative beads (bottom row) showed no detectable signal under these conditions. (C) Time-lapse images in the FITC channel showed fluorescence release from beads irradiated with either 200 ms or 20 ms pulses. Longer exposure resulted in a faster increase in fluorescence intensity, indicating accelerated cleavage kinetics. (D) The fluorescein fluorescence intensity at each imaging time point was converted to estimated concentrations using calibration curves. Under 200 ms exposure, fluorescein concentration reached approximately 60 µM at 60 minutes and exceeded 90 µM at 120 minutes. With 20 ms exposure, the concentration increased gradually and stabilized at around 80 µM after 180 minutes.

Fluorescein-conjugated OBOC beads were suspended at a concentration of 1.5×10^6^ particles/mL in PBST and loaded into nanopens using optimized parameters for import volume, penning algorithm, and Opto-Electropositioning (OEP). To address bead size variability between synthesis lots, the CNN Cell Detect algorithm was selected, which significantly improved the efficiency of transporting individual particles into nanopen using OEP—a process referred to as penning—as well as counting accuracy. To determine the optimal timing for imaging after compound release, we applied the Wilke–Chang equation to estimate diffusion rates^13^. Assuming a compound molecular weight of 400 Da and a nanopen height of 370 µm, the estimated diffusion coefficient was calculated to be 4.23×10^−10^ m^2^/s, suggesting a diffusion time of approximately 2 minutes. Based on this calculation, imaging was initiated at least 2 minutes after UV irradiation, with intervals ranging from 30 minutes to 2 hours depending on the assay (Figure 3B).

To quantify released compounds, carboxyfluorescein solutions were used to generate calibration curves correlating fluorescein fluorescence intensity with concentration and exposure time. Theoretical calculations indicated that complete linker cleavage could yield fluorescein concentrations exceeding 100 µM, but such high levels risk cytotoxicity. A concentration of 10 µM was deemed sufficient for phenotypic changes, and imaging conditions were optimized to visualize concentrations up to 50 µM. Exposure times beyond 750 ms caused overexposure, so 500 ms was selected for FITC imaging. (Figure S9). Validation experiments using synthesized fluorescein-conjugated (positive) and unconjugated (negative) beads confirmed successful compound release. Positive beads exhibited fluorescence within one minute, stabilizing around 15 µM after 10 minutes, while negative beads showed no change. These results demonstrated the feasibility of using OBOC-DEL beads for compound release and subsequent cell-based evaluation. (Figure S11). Interestingly, an unexpected pause in the experiment provided us an additional insight that a single 200 ms UV (390 nm) exposure was sufficient to release compounds, with fluorescein fluorescence detected 20 minutes later. This prompted a reevaluation of exposure duration, showing that shorter exposures could achieve effective cleavage while minimizing phototoxicity. (Figure S12A).

We also investigated the impact of CO_2_ flow on compound retention. Continuous CO_2_ blowing during assays caused fluorescein leakage into adjacent nanopens, limiting accumulation. In contrast, stopping CO₂ flow after initial channel clearing resulted in a fivefold increase in fluorescein concentration—12 µM vs. 63 µM after 60 minutes. This confirmed that halting CO_2_ flow enhances compound retention and concentration within nanopens (Figure S12B, 12D). Further experiments compared 200 ms and 20 ms UV (390 nm) exposures. At 200 ms, fluorescein concentrations reached 20 µM in 30 minutes, 60 µM in 60 minutes, and exceeded 90 µM in 120 minutes. At 20 ms, concentrations rose more gradually, reaching 10 µM in 30 minutes and stabilizing around 80 µM after 180 minutes. These results demonstrated that UV exposure time could be tuned to control compound release kinetics and concentration (Figure 3C, 3D).

Based on these findings, we established a standardized workflow for compound release: single synthesized fluorescein-conjugated are loaded into nanopens, CO_2_ was applied for >30 minutes to remove residual liquid and then stopped, UV (390 nm) was irradiated for 200 ms, and FITC imaging was performed at an exposure time of 500 ms for up to 6 hours. Fluorescein fluorescence intensity was analyzed using Beacon’s Image Analyzer, and compound concentration was estimated via calibration curves. This optimized protocol ensured efficient and reproducible compound release with minimal cellular damage, forming the foundation for downstream phenotypic screening in the Beacon system.

### Verification of the effect of model compounds on cellular phenotype

To validate the Beacon® Optofluidic System for phenotypic screening, we conducted a series of experiments to assess both the viability and responsiveness of engineered reporter cells to model compounds. PC-3 cells stably expressing FKBP12^F36V^-EGFP were used as the reporter system^14^ . Although these cells are naturally adherent, they were adapted for microfluidic loading by detachment and handling in non-adherent conditions. Prior to loading, cells were filtered through a 40 µm strainer and suspended at 1.5×10^6^ cells/mL. Beacon loading parameters—including cage speed, voltage, and frequency—were optimized to improve penning efficiency, and the CNN Watershed algorithm was applied for accurate cell detection. To establish normal cell behavior, cells were loaded into nanopens and monitored for 16 hours without compound exposure. Time-lapse imaging revealed sustained EGFP fluorescence throughout the observation period. Cell proliferation was observed in approximately 13% of nanopens at 6 hours, and by 16 hours, cell doubling was confirmed in approximately 35% of nanopens. EGFP fluorescence intensity remained stable, with no cells exhibiting values below 5,000. Based on these findings, a fluorescence intensity threshold of 5,000 was set for subsequent assays to ensure that only viable cells were included in the analysis (Figure 4A, 4C).

**Figure 4.**
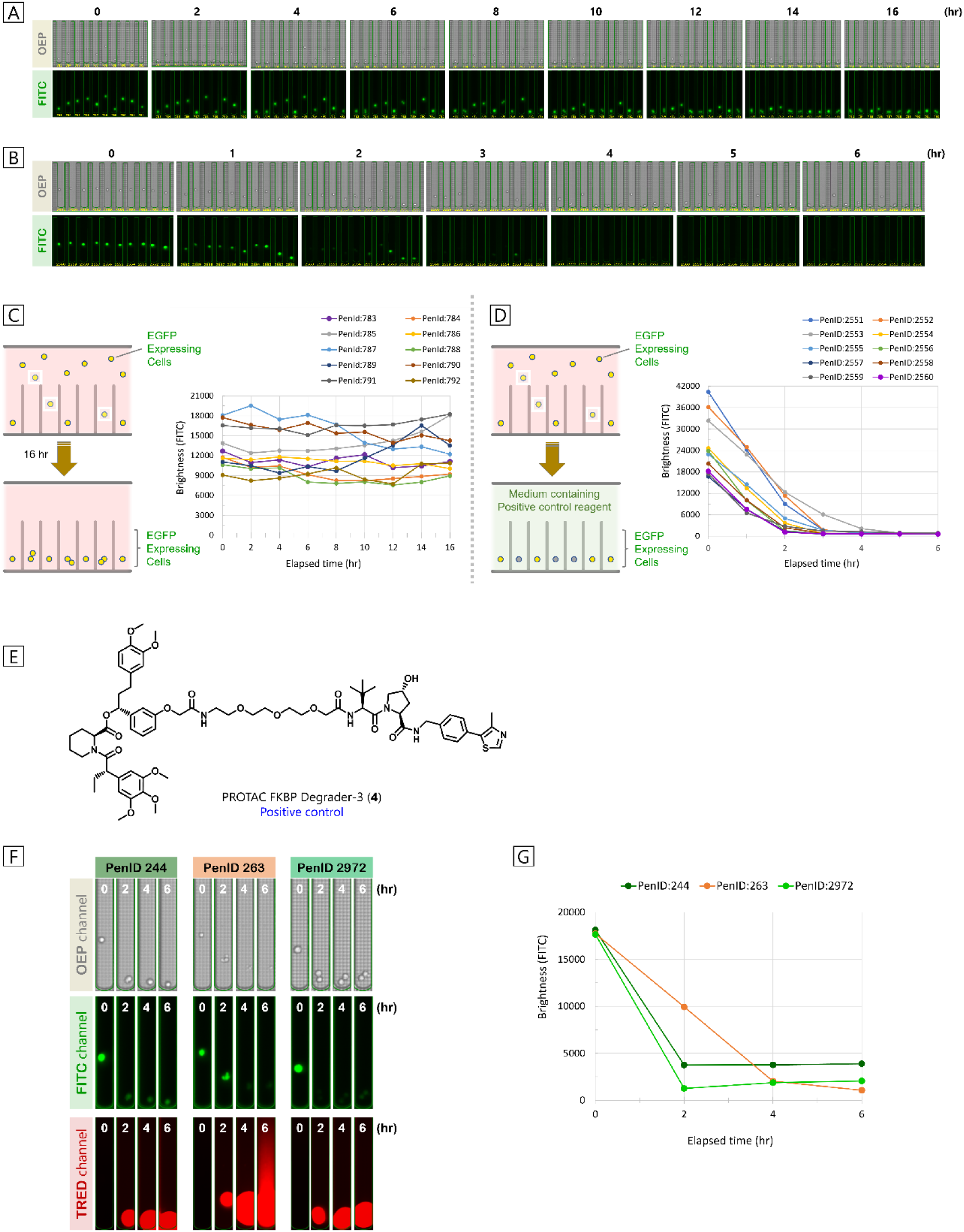
Time-lapse imaging and EGFP fluorescence intensity analysis of single cells on the OptoSelect chip. (A) Single cells were loaded into nanopens on the OptoSelect chip and monitored every 2 hours over a 16-hour period. Time-lapse imaging revealed sustained EGFP fluorescence in most cells throughout the observation. Cell proliferation was observed in approximately 13% of nanopens at 6 hours and in about 35% of nanopens at 16 hours. (B) After cell loading, the medium was replaced with one containing the positive control compound (PROTAC FKBP Degrader-3), and EGFP fluorescence intensity was monitored. Time-lapse imaging showed cell proliferation as early as 2 hours after the start of observation. (C) Quantitative analysis of EGFP fluorescence intensity in individual cells from (A) indicated that the average intensity remained nearly constant, with no significant decrease observed during the 16-hour observation period. (D) Quantitative analysis of individual cells from (B) revealed a marked decrease in EGFP fluorescence, and by 6 hours, fluorescence was undetectable in all cells. (E) Chemical structure of PROTAC FKBP Degrader-3. (F) Although non-specific staining by the viability dye (SYTOX Orange Nucleic Acid Stain) was observed over time, only one nanopen showed widespread fluorescence in the TRED channel after the 6-hour assay (e.g., PenID 263). In addition, bright-field imaging in the OEP channel confirmed that live cells retained clear cellular boundaries without morphological distortion (e.g., PenID 244). Combining these features from both channels allowed clear discrimination between live and dead cells, and overall viability exceeded 99%. Furthermore, cell proliferation was observed in approximately 12% of nanopens (e.g., PenID 2972). (G) During the 6-hour observation period, differences in cellular behavior were observed at the single-cell level, including cells showing either rapid or gradual decreases in EGFP fluorescence intensity.

Next, to verify the system’s ability to detect compound-induced protein degradation, we introduced a model compound—PROTAC FKBP Degrader-3^14,15^— at a final concentration of 1.0 µM (Figure 4E). Cells were cultured for 6 hours, and EGFP fluorescence intensity was monitored every 2 hours. A marked decrease in EGFP fluorescence was observed as early as 2 hours, and by 6 hours, fluorescence was nearly undetectable across all cells. Despite initial variability in EGFP levels, the decline pattern was consistent, confirming robust compound-induced degradation (Figure 4B, 4D).

Having confirmed degradation, we next assessed cytotoxicity by adding SYTOX Orange Nucleic Acid Stain (final concentration, 2.0 µM) after 22 hours of incubation. Weak fluorescence was detected in the TRED channel; however, bright-field imaging revealed that most cells retained well-defined boundaries and exhibited no morphological alterations, suggesting that staining was largely non-specific and membrane integrity remained intact. In contrast, cell death was observed in 2–3 cells per field (1.4–2.0%), characterized by loss of boundaries and diffuse TRED fluorescence. Notably, approximately 43% of nanopens displayed cell proliferation at 22 hours, and combined with a viability exceeding 98%, these findings demonstrate that the compound promotes protein degradation with minimal impact on cell viability (Figure S16).

Based on these findings, the assay conditions were optimized for small-molecule screening. A combined medium containing PROTAC FKBP Degrader-3 and SYTOX Orange was prepared, and cells were cultured for 6 hours following a single UV irradiation (390 nm, 200 ms). Imaging was performed every 2 hours using exposure times of 2,000 ms for the FITC channel and 1,000 ms for the TRED channel. During the assay, cell proliferation was observed in 11–14% of nanopens. At the 6-hour time point, strong SYTOX Orange fluorescence was detected; however, except for a single case, the signal did not spread throughout the nanopen. Bright-field imaging confirmed that most cells maintained clear morphology without distortion. These observations indicate that, despite some non-specific staining, reliable discrimination between live and dead cells was achieved, and overall cell viability exceeded 99% after the assay. This approach enabled single-cell analysis of diverse behaviors, including subtle differences in cell size and protein expression intensity prior to the assay, presence or absence of cell division during the assay, sedimentation patterns within nanopens, and variability in EGFP fluorescence decay rates (Figure 4F, 4G). In light of these preliminary results, the concentration of the viability dye was reduced in subsequent experiments to minimize non-specific staining of live cells and prevent signal saturation in the TRED channel.

Collectively, these results indicate that the Beacon system enables high-resolution phenotypic evaluation of compound activity while minimizing phototoxicity. The engineered PC-3 cells demonstrated stable behavior on the Beacon chip and responded predictably to the model compound. The optimized assay conditions—including cell loading, compound exposure, and imaging protocols—established a reliable platform for evaluating cellular phenotypes in response to small molecules, supporting the use of Beacon for single-cell screening and laying the groundwork for broader application in drug discovery.

### Validation of the single-bead, single-cell co-culture screening workflow

To establish a high-resolution phenotypic screening platform using the Beacon® Optofluidic System, we developed a workflow that enables precise 1:1 co-culture of single OBOC-DEL beads and individual reporter cells within nanoliter-scale nanopens. This approach, central to the PhenoDEL concept, allows direct observation of compound-induced cellular responses and identification of active compounds via DNA barcoding and sequencing. In this study, PROTAC FKBP Degrader-3 photo-linker OBOC-DEL beads were synthesized via solid-phase synthesis and DNA conjugation, incorporating a photo-cleavable linker for compound release upon UV (390 nm) irradiation. Each bead carried a unique compound and DNA barcode, allowing for precise tracking and identification following phenotypic screening (Figure 5A).

**Figure 5.**
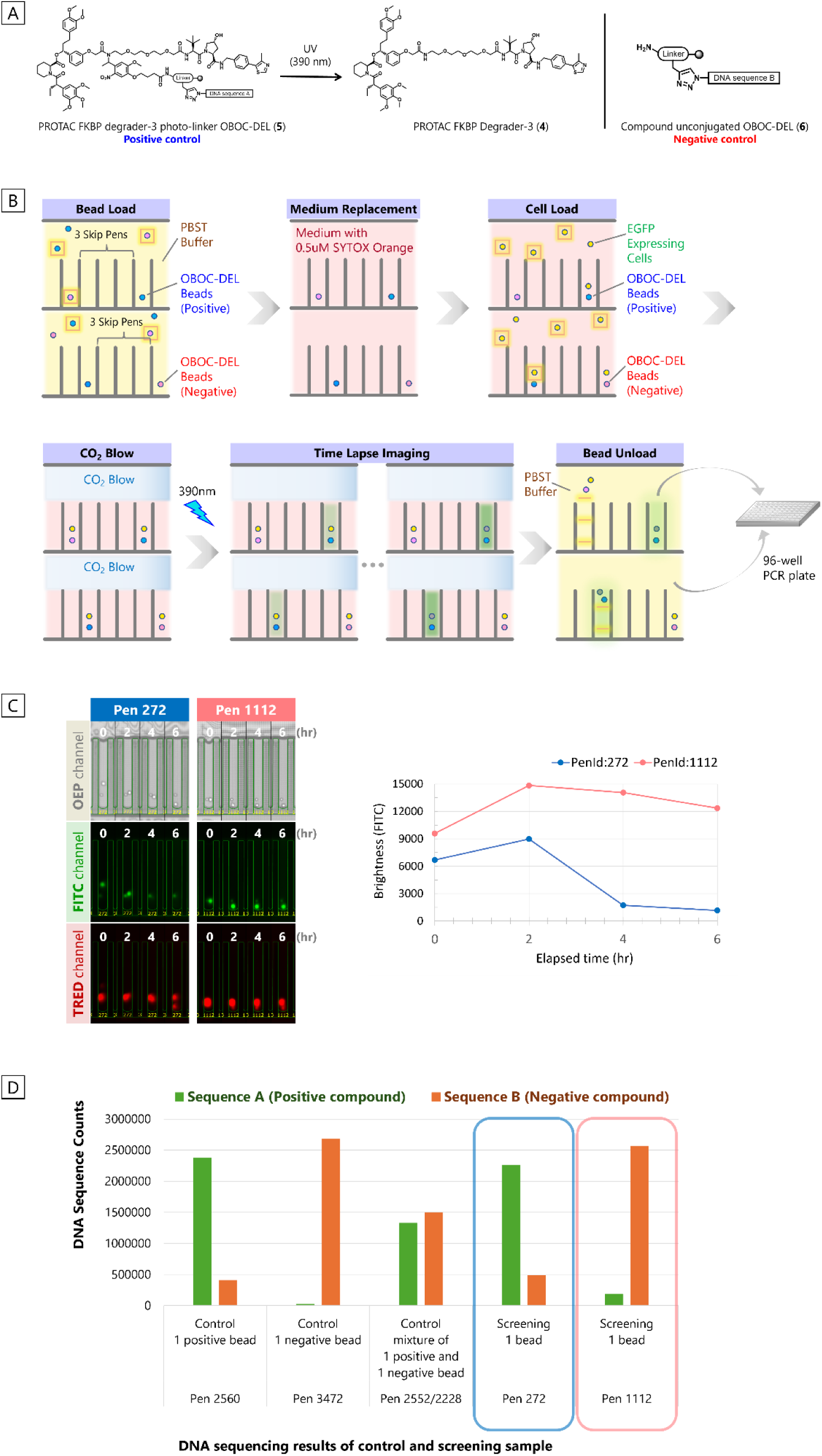
The evaluation of cell based screening using on-bead compounds on the Beacon platform. (A) Chemical structure of the PROTAC FKBP Degrader-3 photo-linker OBOC-DEL bead. The photo-cleavable linker enables compound release upon UV irradiation. Each bead carries a unique compound and a corresponding DNA barcode, allowing for post-screening identification of hit compounds via sequencing. (B) Experimental workflow on the Beacon system. A single cell was loaded into a nanopen containing one OBOC-DEL bead. To prevent compound diffusion outside the nanopen, the medium in the channel was removed. After UV (390 nm) irradiation, cells were monitored for 6 hours. Nanopens containing cells with or without changes in EGFP fluorescence intensity were selected, and beads from these pens were recovered for NGS analysis. (C) Phenotypic screening of 1:1 OBOC-DEL bead–cell co-cultures using Beacon. Nanopens containing one bead and one cell were imaged every two hours after UV (390 nm) irradiation. Most cells exhibited clear boundaries (OEP) and were not stained by the nucleic acid staining reagent (TRED), indicating stable survival within the nanopen. In some cells, EGFP fluorescence intensity decreased, while others showed little to no change even after 6 hours. After 6 hours of culture, PenID 272 showed a decrease in brightness, whereas PenID 1112 exhibited no notable change.

Within the Beacon workflow, individual OBOC-DEL beads were loaded into nanopens, followed by single FKBP12^F36V^-EGFP reporter cells. After confirming successful penning of both bead and cell, UV (390 nm) irradiation (200 ms) was applied to cleave the compound from the bead. The released compound diffused into the nanopen and interacted with the co-cultured cell. Time-lapse imaging was performed every two hours over a six-hour period to monitor changes in EGFP fluorescence, serving as a readout for compound activity. Images acquired during the assay were analyzed using Beacon’s image analysis software to extract data on cellular morphology and EGFP fluorescence intensity. Beads from nanopens showing significant phenotypic changes—defined as a EGFP fluorescence intensity decrease of more than 5,000 units—were recovered and transferred to a 96-well plate. The DNA barcode region of each bead was amplified by PCR and subjected to NGS to identify the compound responsible for the observed effect (Figure 5B).

To ensure objective classification, only nanopens containing one bead and one viable cell with clearly detectable EGFP expression before the assay were included in the analysis. Viable cells were defined as those showing distinct boundaries without deformation in the OEP channel and lacking fluorescence diffusion in the TRED channel. This approach confirmed that changes in EGFP fluorescence were caused by compound activity, not by cell damage (Figure S19).

During the screening, most cells remained viable and showed no signs of damage. In FITC channel imaging, some cells exhibited a marked decrease in EGFP fluorescence, while others showed minimal change. This variability may suggest differential activity of compounds encoded on the beads. In the TRED channel, fluorescence signals were observed on the OBOC-DEL beads due to SYTOX staining of DNA barcodes present on the bead surface. However, these signals were distinguishable from cellular fluorescence based on differences in size and circularity, allowing clear discrimination from cells. To correlate phenotypic outcomes with compound identity, beads were recovered from nanopens representing both positive and negative responses and analyzed by NGS (Figure 5C). To validate the specificity of the assay, control experiments were performed in which OBOC-DEL beads were loaded and unloaded without co-culturing with cells. DNA barcodes from these beads were sequenced to establish baseline profiles. The analyzed beads included those with compounds covalently attached via photo-cleavable linkers (positive beads), beads without any compound conjugation (negative beads), and a mixed population containing both types at a 1:1 ratio. Recovered beads were subjected to two rounds of PCR amplification, and the resulting products were sequenced using Illumina high-throughput sequencing. The data were processed and analyzed to determine enrichment patterns and identify active compounds. Comparative analysis of sequencing data from control and test samples confirmed the specificity and reproducibility of the screening workflow (Figure 5D).

This integrated approach—combining compound release, phenotypic tracking, and molecular decoding—demonstrates the power of Beacon-based PhenoDEL screening. It enables high-resolution, activity-based screening of DNA-encoded libraries at the single-cell level, overcoming limitations of conventional droplet-based methods such as lack of spatial control, inability to track individual cell behavior, and difficulty in eliminating false positives. By enabling precise 1:1 pairing of beads and cells, real-time tracking of cellular responses, and direct linkage between phenotype and compound identity, this workflow supports scalable and mechanism-driven drug discovery. It is compatible with various cell types and reporter systems, making it a versatile platform for functional genomics and small molecule screening.

## Discussion(Summary)

In this study, we developed and named a novel screening strategy, PhenoDEL (Phenotypic DNA-Encoded Library), which integrates OBOC-DEL with the Beacon® optofluidic system to enable phenotype-driven identification of small molecules that induce target protein degradation. This platform allows for high-resolution, activity-based screening at the single-cell level by co-culturing individual beads and cells within nanoliter-scale NanoPen™ chambers.

PhenoDEL offers several key advantages over conventional droplet-based and affinity-based screening methods. The Beacon system enables precise 1:1 loading of beads and cells into nanopens, which can be visually confirmed through real-time imaging. Importantly, the single-cell resolution of PhenoDEL allows for rigorous quality control by excluding dead or damaged cells from analysis, which is not feasible in pooled cell screening. As a result, the platform provides more accurate and reliable phenotypic data. This spatial control allows for the automatic acquisition of fluorescence intensity both before and after compound elution—an approach not feasible in traditional screening formats. By capturing dynamic phenotypic changes rather than relying solely on endpoint measurements, PhenoDEL minimizes false positives and enhances assay reliability.

We demonstrated that compound release from OBOC-DEL beads can be finely tuned by adjusting UV (390 nm) irradiation time. A brief 200 ms exposure was sufficient to achieve biologically relevant compound concentrations without compromising cell viability. Furthermore, halting CO₂ flow during the assay significantly improved compound retention within the nanopen, enhancing sensitivity and reproducibility. The modularity of PhenoDEL supports high-throughput screening of large chemical libraries and is compatible with various cell types and reporter systems. Importantly, the platform is not restricted to specific cell types^16^, allowing broad applicability across diverse biological models. We have successfully applied the Beacon system to a variety of cell types, including suspension cells (e.g., CHO-S), adherent cells (e.g., HEK-293), and hybridoma cells (e.g., OKT3). These applications demonstrated consistent operability and analytical precision across different biological contexts. To support scalable screening, the Beacon system enables simultaneous operation of up to four chips. Among them, the OptoSelect® 20k chip (Bruker Cellular Analysis, #750-00019) contains approximately 20,000 nanopens, allowing for the screening of up to 80,000 unique samples in a single run. In our proof-of-concept study using FKBP12^F36V^-EGFP reporter cells, we successfully identified compounds that induced target protein degradation, validating the platform’s utility for PROTAC discovery. Furthermore, although additional validation is warranted, our findings suggest that this method may be broadly applicable to various cell types for phenotypic evaluation, as even inherently adherent cells were successfully handled in a suspended-like manner on the chip under the current experimental conditions. A key feature of the PhenoDEL workflow is the ability to recover beads from individual nanopens and decode their DNA barcodes via PCR and NGS. This enables a direct and unambiguous linkage between compound identity and cellular phenotype, facilitating high-resolution, activity-based screening at the single-cell level.

Looking forward, PhenoDEL holds significant potential for expanding the scope of functional small molecule discovery. Its compatibility with diverse biological models—including patient-derived cells, primary cells, and organoids—makes it a promising tool for personalized medicine and disease-specific drug screening. Integration with transcriptomic or proteomic profiling could provide deeper mechanistic insights, while the incorporation of machine learning–based image analysis may further enhance the efficiency and precision of hit identification. Moreover, PhenoDEL’s ability to track real-time cellular responses opens new avenues for studying dynamic biological processes, such as protein degradation kinetics, pathway modulation, and cellular heterogeneity. These capabilities position PhenoDEL as a powerful platform for mechanism-driven drug discovery, particularly in the identification of novel PROTACs, molecular glue degraders, and other proximity-inducing molecules. As future considerations, further optimization of experimental conditions will be required for scaling up the workflow, and preliminary validation of single-cell on-chip culture compatibility should be conducted when applying the system to different cell types.

In conclusion, PhenoDEL represents a significant advancement in DNA-encoded library screening technology. By overcoming the limitations of existing methods and enabling high-resolution, activity-based screening at the single-cell level, it provides a scalable and versatile framework for next-generation drug discovery and functional genomics.

## Acknowledgements

We gratefully acknowledge the experimental support and insightful discussions provided by our colleagues at Mitsubishi Tanabe Pharma Corporation and Nikon Corporation. Their contributions were essential to the progress of this research. We also extend our appreciation to the team at Bruker Cellular Analysis (formerly known as Berkeley Lights Inc.) for their expert technical advice and assistance.

## Author contributions

Yuichi Onda and Yurika Ochi contributed equally. Yuichi Onda, Yurika Ochi and T. A. conceived and designed the experiments. Yuichi Onda, S.T. and Y.T. performed chemical synthesis. T. A., M.K. and K.Y. prepared and engineered cell lines. Yurika Ochi, Yuichi Onda, T. A., and M.K. conducted Beacon experiments and data analysis. K.O., M.T., D.S., M.K. Y.T. and T.U. provided advice and helped with data analysis. Yuichi Onda and Yurika Ochi wrote the original draft. The manuscript was written with contributions from all authors and was corrected by all authors.

## Data availability

All data supporting the findings of this study are available within the article and Supplementary Information.

## References

1 a) Chamberlain, P. P., Hamann, L. G. Development of targeted protein degradation therapeutics. Nature Chemical Biology 15, 937–944 (2019). 10.1038/s41589-019-0362-y b) Tsai, J. M., Nowak, R. P., Ebert, B. L., Fischer, E. S. Targeted protein degradation: from mechanisms to clinic. Nature Reviews Molecular Cell Biology (2024). 10.1038/s41580-024-00729-9 c) Zhong, G., Chang, X., Xie, W., Zhou, X. Targeted protein degradation: advances in drug discovery and clinical practice. Signal Transduction and Targeted Therapy 9, 308 (2024). 10.1038/s41392-024-02004-x

2 a) Liu, X., Ciulli, A. Proximity-Based Modalities for Biology and Medicine. ACS Central Science 9, 1269–1284 (2023). 10.1021/acscentsci.3c00395 b) Singh, S., Tian, W., Severance, Z. C., Chaudhary, S. K., Anokhina, V., Mondal, B., Pergu, R., Singh, P., Dhawa, U., Singha, S., Choudhary, A. Proximity-inducing modalities: the past, present, and future. Chemical Society Reviews 52, 5485–5515 (2023). 10.1039/d2cs00943a

3 a) Dong, G., Ding, Y., He, S., Sheng, C. Molecular Glues for Targeted Protein Degradation: From Serendipity to Rational Discovery. Journal of Medicinal Chemistry 64, 10606–10620 (2021). 10.1021/acs.jmedchem.1c00895 b) Holdgate, G. A., Bardelle, C., Berry, S. K., Lanne, A., Cuomo, M. E. Screening for molecular glues – Challenges and opportunities. SLAS Discovery 29, 100136 (2024). 10.1016/j.slasd.2023.12.008 c) Domostegui, A., Nieto-Barrado, L., Perez-Lopez, C., Mayor-Ruiz, C. Chasing molecular glue degraders: screening approaches. Chemical Society Reviews 51, 5498–5517 (2022). 10.1039/d2cs00197g d) Darling, S., Bajrami, I., West, S. C. From serendipity to strategy: molecular glue degraders in cancer therapeutics. Critical Reviews in Biochemistry and Molecular Biology (2025). 10.1080/10409238.2025.2564068

4 a) Neri, D., Lerner, R. A. DNA-Encoded Chemical Libraries: A Selection System Based on Endowing Organic Compounds with Amplifiable Information. Annu. Rev. Biochem. 87, 479–502 (2018). 10.1146/annurev-biochem-062917-012550 b) Favalli, N., Bassi, G., Scheuermann, J., Neri, D. DNA-encoded chemical libraries – achievements and remaining challenges. FEBS Letters 592, 2168–2180 (2018). 10.1002/1873-3468.13068 c) Goodnow, R. A. Jr., Dumelin, C. E., Keefe, A. D. DNA-encoded chemistry: enabling the deeper sampling of chemical space. Nature Reviews Drug Discovery 16, 131–147 (2017). 10.1038/nrd.2016.213 d) Gironda-Martínez, A., Donckele, E. J., Samain, F., Neri, D. DNA-Encoded Chemical Libraries: A Comprehensive Review with Succesful Stories and Future Challenges. ACS Pharmacol. Transl. Sci. 4, 1265–1279 (2021). 10.1021/acsptsci.1c00118 e) Valeur, E., Jimonet, P. New Modalities, Technologies, and Partnerships in Probe and Lead Generation: Enabling a Mode-of-Action Centric Paradigm. Journal of Medicinal Chemistry 61, 9004–9029 (2018). 10.1021/acs.jmedchem.8b00378

5 a) Disch, J. S., Duffy, J. M., Lee, E. C. Y., Gikunju, D., Chan, B., Levin, B., Monteiro, M. I., Talcott, S. A., Lau, A. C., Zhou, F., Kozhushnyan, A., Westlund, N. E., Mullins, P. B., Yu, Y., von Rechenberg, M., Zhang, J., Arnautova, Y. A., Liu, Y., Zhang, Y., McRiner, A. J., Keefe, A. D., Kohlmann, A., Clark, M. A., Cuozzo, J. W., Huguet, C., Arora, S. Bispecific Estrogen Receptor α Degraders Incorporating Novel Binders Identified Using DNA-Encoded Chemical Library Screening. Journal of Medicinal Chemistry 64, 5049–5066 (2021). 10.1021/acs.jmedchem.1c00127 b) Chen, Q., Liu, C., Wang, W., Meng, X., Cheng, X., Li, X., Cai, L., Luo, L., He, X., Qu, H., Luo, J., Wei, H., Gao, S., Liu, G., Wan, J., Israel, D. I., Li, J., Dou, D. Optimization of PROTAC Ternary Complex Using DNA Encoded Library Approach. ACS Chemical Biology 18, 25–33 (2023). 10.1021/acschembio.2c00797 c) Mason, J. W., Chow, Y. T., Hudson, L., Tutter, A., Michaud, G., Westphal, M. V., Shu, W., Ma, X., Tan, Z. Y., Coley, C. W., Clemons, P. A., Bonazzi, S., Berst, F., Briner, K., Liu, S., Zécri, F. J., Schreiber, S. L. DNA-encoded library-enabled discovery of proximity-inducing small molecules. Nature Chemical Biology 20, 170–179 (2024). 10.1038/s41589-023-01458-4 d) Liu, S., Tong, B., Mason, J. W., Ostrem, J. M., Tutter, A., Hua, B. K., Tang, S. A., Bonazzi, S., Briner, K., Berst, F., Zécri, F. J., Schreiber, S. L. Rational Screening for Cooperativity in Small-Molecule Inducers of Protein–Protein Associations. Journal of the American Chemical Society 145, 23281–23291 (2023). 10.1021/jacs.3c08307

6 a) MacConnell, A. B., McEnaney, P. J., Cavett, V. J., Paegel, B. M. DNA-encoded solid-phase synthesis: encoding language design and complex oligomer library synthesis. ACS Comb. Sci. 17, 518–534 (2015). 10.1021/acscombsci.5b00106 b) Mendes, K. R., Malone, M. L., Ndungu, J. M., Suponitsky-Kroyter, I., Cavett, V. J., McEnaney, P. J., MacConnell, A. B., Doran, T. M., Ronacher, K., Stanley, K., Utset, O., Walzl, G., Paegel, B. M., Kodadek, T. High-throughput identification of DNA-encoded IgG ligands that distinguish active and latent Mycobacterium tuberculosis infections. ACS Chem. Biol. 12, 234–243 (2017). 10.1021/acschembio.6b00855 c) Cochrane, W. G., Malone, M. L., Dang, V. Q., Cavett, V., Satz, A. L., Paegel, B. M. Activity-based DNA-encoded library screening. ACS Comb. Sci. 21, 425–435 (2019). 10.1021/acscombsci.9b00037 d) Dixit, A., Paegel, B. M. Solid-phase DNA-encoded library synthesis: a master builder’s instructions. Nat. Protoc. 19, 1–40 (2025). 10.1038/s41596-025-01190-4

7 a) Barhoosh, H., Dixit, A., Cochrane, W. G., Cavett, V., Prince, R. N., Blair, B. O., Ward, F. R., McClure, K. F., Patten, P. A., Paulick, M. G., Paegel, B. M. Activity-Based DNA-Encoded Library Screening for Selective Inhibitors of Eukaryotic Translation. ACS Cent. Sci. 10, 1960–1968 (2024). 10.1021/acscentsci.4c01218

8 a) Bushman, J. W., Deng, W., Samarasinghe, K. T. G., Liu, H.-Y., Li, S., Vaish, A., Golkar, A., Ou, S.-C., Ahn, J., Harijan, R. K., Han, H., Duffy, T., Imler, E., Wu, J., Rawson, R., Liu, L., Ndungu, J. M., Hocker, M. D., Roy, A., Senese, A. D., Lin, C.-C., Tran, D., Campos, A., Blanco, G., Toth, J. I., Velentza, A., Burrit, A., Parker, G. S., Thompson, P. A., Bailey, S., den Besten, W., Voss, S., Zhou, B., Ashton, K. S., Bedel, O., Min, J., Pots, P. R. Discovery of a VHL molecular glue degrader of GEMIN3 by Picowell RNA-seq. bioRxiv (2025). 10.1101/2025.03.19.644003

9 a) Mocciaro, A., Roth, T. L., Bennett, H. M., Soumillon, M., Shah, A., Hiatt, J., Chapman, K., Marson, A. & Lavieu, G. Light-activated cell identification and sorting (LACIS) for selection of edited clones on a nanofluidic device. *Commun*. Biol. 1, 41 (2018). 10.1038/s42003-018-0034-6 b) Jorgolli, M., Nevill, T., Winters, A., Chen, I., Chong, S., Lin, F.-F., Mock, M., Chen, C., Le, K., Tan, C., Jess, P., Xu, H., Hamburger, A., Stevens, J., Munro, T., Wu, M., Tagari, P., Miranda, L. P. Nanoscale integration of single cell biologics discovery processes using optofluidic manipulation and monitoring. Biotechnol. Bioeng. 116, 2393–2411 (2019). 10.1002/bit.27024

10 a) Le, K., Tan, C., Le, H., Tat, J., Zasadzinska, E., Diep, J., Zastrow, R., Chen, C., Stevens, J. Assuring clonality on the Beacon Digital Cell Line Development Platform. Biotechnol. J. 15, 1900247 (2020). 10.1002/biot.201900247

11 a) Winters, A., McFadden, K., Bergen, J., Landas, J., Berry, K. A., Gonzalez, A., Salimi-Moosavi, H., Murawsky, C. M., Tagari, P., King, C. T. Rapid single B cell antibody discovery using nanopens and structured light. mAbs 11, 1025–1035 (2019). 10.1080/19420862.2019.1624126

12 a) Xiong, T., Wang, G., Yu, P., Li, Z., Li, D., Zhang, J., Lu, S., Yang, R., Lian, X., Mi, J., Ma, R., Li, Z., Marcucci, G., Zhao, T., Caligiuri, M. A., Yu, J. CAR-T cells targeting CD155 reduce tumor burden in preclinical models of leukemia and solid tumors. J. Clin. Invest. 135, e189920 (2025). 10.1172/JCI189920

13 a) Wilke, C. R., Chang, P. Correlation of diffusion coefficients in dilute solutions. A.I.Ch.E. J. 1, 264–270 (1955). 10.1002/aic.690010222

14 a) Ottis, P., Toure, M., Cromm, P. M., Ko, E., Gustafson, J. L., Crews, C. M. Assessing different E3 ligases for small molecule induced protein ubiquitination and degradation. ACS Chem. Biol. 12, 2570–2578 (2017). 10.1021/acschembio.7b00485

15 a) Bradner, J., Roberts, J., Nabet, B., Winter, G., Phillips, A. J., Heffernan, T. P., Buckley, D. Targeted protein degradation to attenuate adoptive T-cell therapy associated adverse inflammatory responses. **WO2017/024318 A1** (2017).

16 a) Le, K., Tan, C., Le, H., Tat, J., Zasadzinska, E., Diep, J., Zastrow, R., Chen, C. & Stevens, J. Assuring clonality on the Beacon Digital Cell Line Development Platform. Biotechnol. J. 15, 1900247 (2020). 10.1002/biot.201900247 b) Zhang, Y., Park, M., Ghoda, L. Y., Zhao, D., Vaero, M., Nafe, E., Gonzalez, A., Ly, K., Parcutea, B., Cho, H., Gong, X., Chen, F., Harada, K., Chen, Z., Nguyen, L. X. T., Pchorri, F., Chen, J., Song, J., Forman, S. J., Amanam, I., Zhang, B., Jin, J., Wams, J. C. & Marcucci, G. IL1RAP-specific T cell engager depletes acute myeloid leukemia stem cells. J. Hematol. Oncol. 17, 67 (2024). 10.1186/s13045-024-01586-x c) Colina, A. S., Shah, V., Shah, R. K., Kozlik, T., Dash, R. K., Terhune, S. & Zamora, A. E. Current advances in experimental and computational approaches to enhance CAR T cell manufacturing protocols and improve clinical efficacy. Front. Mol. Med. 4, 1310002 (2024). 10.3389/fmmed.2024.1310002 d) Rienzo, M., Lin, K.-C., Mobilia, K. C., Sackmann, E. K., Kurz, V., Navidi, A. H., King, J., Onorato, R. M., Chao, L. K., Wu, T., Jiang, H., Valley, J. K., Lionberger, T. A. & Leavell, M. D. High-throughput optofluidic screening for improved microbial cell factories via real-time micron-scale productivity monitoring. Lab Chip 21, 2901–2912 (2021). 10.1039/d1lc00389e

