## Supplementary information for "PhenoDEL as a Novel Screening Strategy Based on Intracellular Protein Degradation Activity"

### 1. Abbreviations

| | |
|---|---|
| Beacon | : Beacon® Optofluidic System |
| Brine | : saturated sodium chloride solution |
| Chip | : OptoSelect™ chip |
| DAPI | : 4’,6-diamidine-2’-phenylindole dihydrochloride |
| DCM | : dichloromethane |
| DIPEA | : N, N’-diisopropylethylamine |
| DMF | : N, N’-dimethylformamide |
| DMSO | : dimethyl sulfoxide |
| DEL | : DNA encoded library |
| EtOAc | : ethyl acetate |
| ESI | : electrospray ionization |
| FKBP | : FK506-binding protein |
| FK506 | : Tacrolimus, CAS 104987-11-3 |
| Fmoc | : 9-fluorenylmethyloxycarbonyl |
| FOV | : Field of View |
| HATU | :1-[bis(dimethylamino)methylene]-1H-1,2,3-triazolo[4,5b] pyridinium 3-oxide hexafluorophosphate |
| HPLC | : high performance liquid chromatography |
| Nanopen | : NanoPen™ chambers |
| ms | : millisecond(s) |
| OEP | : Opto-Electropositioning |
| OBOC | : one bead one compound |
| OBOC-DEL | : one bead one compound DNA encoded library |
| Photo-Linker | : 4-[4-(1-aminoethyl)-2-methoxy-5-nitrophenoxy]butanoic acid |
| PROTAC | : proteolysis targeting chimera |
| PROTAC FKBP Degrader-3 | : CAS 2079056-43-0 |
| SPPS | : solid phase peptide synthesis |
| TFA | : trifluoroacetic acid |
| TPS | : Target and Pen Selection |
| UPLCMS | : ultra performance liquid chromatography mass spectrometry |

### 2. Materials and General Methods

Unless otherwise noted, all reagents and solvents were purchased from commercial sources and used as received. Oligonucleotides were purchased from DNA Technology (Denmark) and IBA (Germany). All DNA sequences are written from 5'- to 3'- end. Carboxylic acids and terminal alkynes were purchased from ABCR, ChemBridge, Sigma-Aldrich, TCI Europe, Alfa Aesar, Matrix Scientific, Enamine Store and Acros Organics. H2O was purified with a Millipore Milli-Q system (Merck). All gel images were captured by a Bio-Rad Chemidoc image system.

Analyses for the compounds: UPLCMS analyses were performed on a BEH C18 2.1 × 50 mm column at a flow rate of 0.6 mL/min with gradient: 5% to 98% B (0 to 1 minutes), 98% B (1 to 1.2 minutes), (A= 0.05% formic acid in water, B= $CH_3CN$ with 0.05% formic acid), at 40 ºC.

Preparations for the compounds: HPLC preparation were performed on a Xbridge C18 (Φ30mm x 50mm, 5μm) column at a flow rate of 50 ml/min with gradient: 50% B (0 to 2 minutes), 50% to 80% (2 to 5 minutes), 80% to 95% (5 to 5.1 minutes), 95% to 100% (5.1 to 6.6 minutes), 100% B (6.6 to 7.5 minutes) (buffer A = 0.05%TFA in mQ millipore water, B = 0.05%TFA in $CH_3CN$) at room temperature.

Beacon® Optofluidic System: The Beacon® Optofluidic System (#110-08004 /Bruker Corporation) uses a core technology called Optofluidics, which is a combination of optics and nanofluidics. Each OptoSelect™ chip (Chip) contains a multitude of NanoPen™ chambers (Nanopen), ranging from thousands to tens of thousands, which function as an alternative to the wells found in a microplate. Cells and other microparticles are transported to each nanopen by Opto-Electropositioning (OEP). After this process, brightfield and fluorescence images are continuously captured, and a multitude of unique assays are performed on the Beacon to characterize single cells. Representing workflows offered by the Beacon include cell line development, antibody discovery, T-cell profiling and cytotoxicity assay. In these and other R&D areas, the Beacon system can dramatically advance our research, saving us significant time, cost, and effort in the race to find critical cells.

### 3. Cell preparation

#### 3.1. Generation of PC-3 cells stably expressing FKBP12$^{F36V}$-EGFP

The plasmid pZFN-AAVS1, which encodes AAVS1-specific ZFNs[1], was constructed by inserting synthesized DNA into a CMV promoter vector. The FKBP12$^{F36V}$-EGFP targeting vector, pZDAP-PDGH1, was constructed by sub-cloning synthesized mPGK promoter- FKBP12$^{F36V}$-EGFP gene into pZDonor-AAVS1 vector (Sigma, St. Louis, MO, USA). PC-3 cells were obtained from ATCC and cultured in F-12K medium (Thermo Fisher Scientific) with 10% fetal bovine serum (Sigma-Aldrich) and 1× Penicillin-Streptomycin solution (FUJI FILM). PC-3 cells expressing FKBP12$^{F36V}$-EGFP were established by co-transfecting pZFN-AAVS1 and pZDAP-PDGH1, using Lipofectamine3000 (Thermo Fisher Scientific). To generate stable cell lines, 0.25 μg/mL Puromycin (InvivoGen) was added to the culture medium 24 hours after transfection. The cells were then cultured at 37°C with 5% $CO_2$ for 2 weeks, cloned, and used for subsequent assays. Established cells were passaged once every 2-3 days.

#### 3.2. Confirmation of activity of degrader-3 by flow cytometry

PC-3 cells stably expressing FKBP12$^{F36V}$-EGFP were cultured in a 150 mm dish (AS ONE) until further use. The cells were washed with PBS (nacalai tesque) and then dissociated using Accutase (Innovative Cell Technologies). The cells were suspended in culture medium and seeded in a 384-well plate (Corning). Degrader-3 (MedChemExpress), diluted in culture medium, was then added to the wells at final concentrations of 0, 1, 3, 10, 30, 100, 300, and 1,000 nM, and incubated for 6 hours. After incubation, the cells were rinsed twice with flow cytometry staining buffer (Thermo Fisher Scientific), resuspended and analyzed using a flow cytometer (iQue3, Sartorius). The median fluorescence intensity of EGFP decreasedin a dose-dependent manner upon treatment with the compound, and the $DC_{50}$ was determined to be 44 nM.

a) Flow cytometric analysis    b) Determination of $DC_{50}$ value

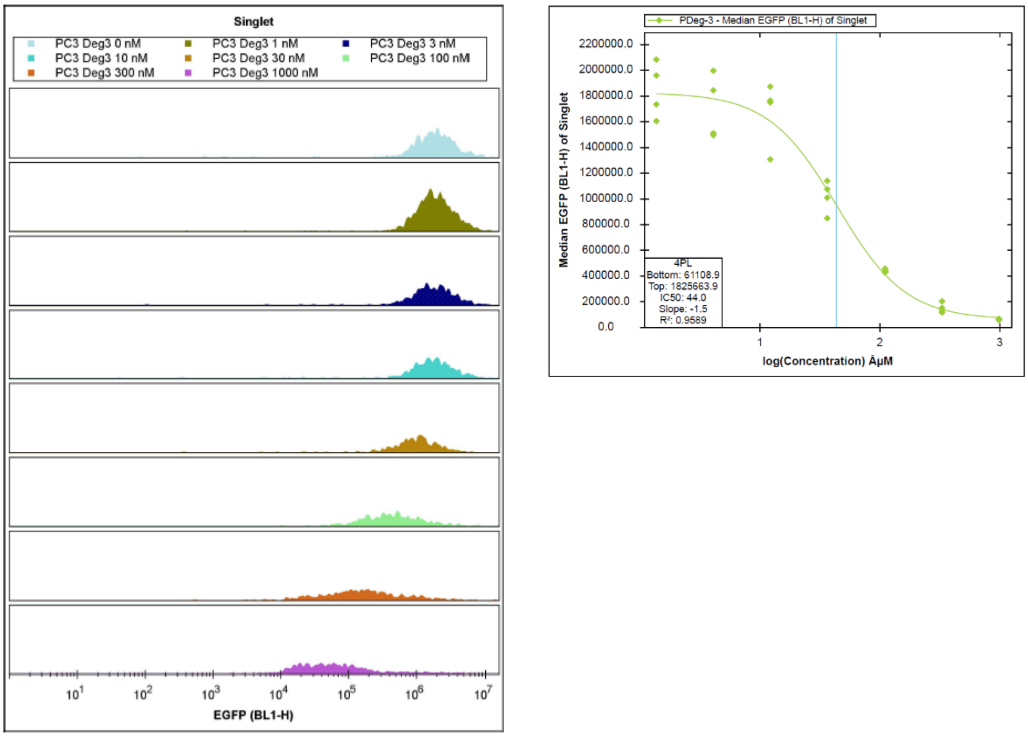

Figure S1. a) Flow cytometric analysis showing FKBP12$^{F36V}$-EGFP degradation in PC-3 cells treated with the PROTAC FKBP Degrader-3. b) Dose-response curve and $DC_{50}$ value determination for FKBP12$^{F36V}$-EGFP degradation in PC-3 cells.

### 4. Compounds synthesis

#### 4-1. Synthesis of fluorescein photo-linker OBOC (1)

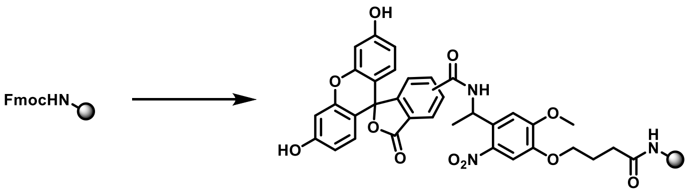

TentaGel® M RAM (91 mg, 20 μmol, 0.22 mmol/g) was swollen with DMF (1.0 mL) in a 10 mL fritted syringe. After 1 hour, the resin was washed with DMF (×1) and treated with 20% piperidine/DMF (1.0 mL) at room temperature. After 1 h, the resin was washed with DMF (×5). The resin was swollen with DMF (1.0 mL) for 10 min. Fmoc-Photo-Linker (42 mg, 80 μmol, 4 equiv.), DIPEA (21 mg, 160 μmol, 8 equiv.) and HATU (30 mg, 80 μmol, 4 equiv.) were added and the mixture was shaken at room temperature. After 2 h, the resin was washed with DMF (×5) and treated with 20% piperidine/ DMF (1.0 mL) at room temperature. After 1 h, the resin was washed with DMF (×5). The resin was swollen with DMF (1.0 mL) for 10 min. 5(6)-carboxyfluorescein (42 mg, 30 μmol, 4 equiv.), DIPEA (21 mg, 160 μmol, 8 equiv.) and HATU (30 mg, 80 μmol, 4 equiv.) were added and the mixture was shaken at room temperature. After 2 h, the resin was washed with DMF (×5) to afford Fluorescein Photo-Linker OBOC (**1**) (91 mg, 20 μmol, quant.).

#### 4-2. Photo-cleavage reaction of fluorescein photo-linker OBOC (1) to yield fluorescein amide (2)

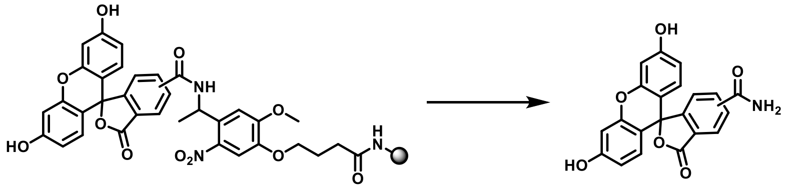

Fluorescein photo-linker OBOC (**1**) (0.1 mg, 0.022 μmol) was swollen with DMF (50 μL) in a 1.5 mL Eppendorf tube®. The Eppendorf tube® was irradiated in a well-ventilated photochemical reactor (EvoluChem PhotoRedOx Box™ and 365 nm EvoluChem™ LED, HepatoChem, MA 01915 USA) at 365 nm at room temperature. After 10 minutes, the resin was filtered, and the filtrate was analyzed via UPLCMS to detect fluorescein amide (**2**). Fluorescein amide (**2**): LC retention time: 0.64 min, MS characterization (*m/z*, $C_{21}H_{13}NO_6$, ESI): calculated [M+H]+: 376.0816, found: 376.0433.

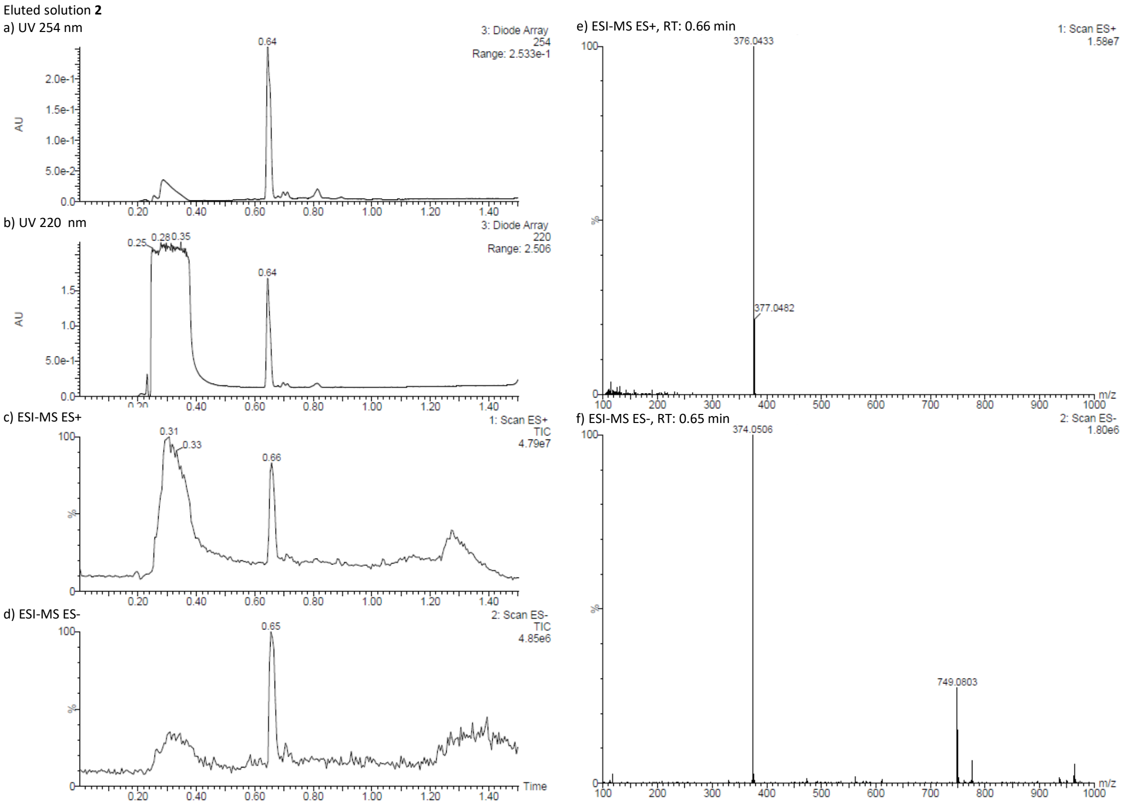

Figure S2. UPLC chromatograms of eluted solution **2**. a) Detection by absorbance at 254 nm, b) Detection by absorbance at 220 nm, c) Scan by ESI-MS ES+, d) Scan by ESI-MS ES-, e) ESI-MS ES+ at 0.66 min, f) ESI-MS ES- at 0.65 min

### 4.3. Synthesis of allyl ester (7)

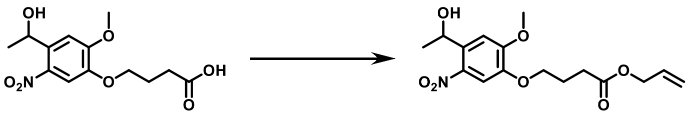

To a solution of 4-[4-(1-hydroxyethyl)-2-methoxy-5-nitro-phenoxy]butanoic acid (1.02 g, 3.41 mmol, 1.00 equiv.) in $CH_3CN$ (2 mL) and DMF (8 mL) was added $Na_2CO_3$ (722 mg, 6.81 mmol, 2.00 equiv.) and 3-bromoprop-1-ene (0.32 mL, 3.7 mmol, 1.10 equiv.) at room temperature. The reaction mixture was stirred at room temperature. (yellow suspension) for 2 hours. To the reaction mixture was added $K_2CO_3$ (471 mg, 3.41 mmol, 1.00 equiv.) and 3-bromoprop-1-ene (0.32 mL, 3.7 mmol, 1.10 equiv.) at room temperature. The whole mixture was stirred at room temperature overnight (yellow suspension). The reaction mixture was concentrated under reduced pressure to remove DMF. To the residue was added $H_2O$, then extracted with EtOAc twice. The combined organic layer was washed with brine, fried over $Na_2SO_4$, filtered and the filtrate was concentrated under reduced pressure to give a crude yellow oil. To the residue was added a mixture of hexane and EtOAc (4/1) to afford pale yellow precipitation. The solid was collected by suction filtration, washed with a mixture of hexane and

EtOAc, dried under reduced pressure to give allyl 4-[4-(1-hydroxyethyl)-2-methoxy-5-nitro-phenoxy]butanoate (**7**) (1.11 g, 3.27 mmol, 96.0% yield) as a pale yellow solid. The residue was used for the next reaction without further purification. Allyl ester (**7**): LC retention time: 0.88 min, MS characterization (*m/z*, $C_{16}H_{21}NO_7$, ESI): calculated [M-$H_2O$+H]+: 322.1286, found: 321.9285.

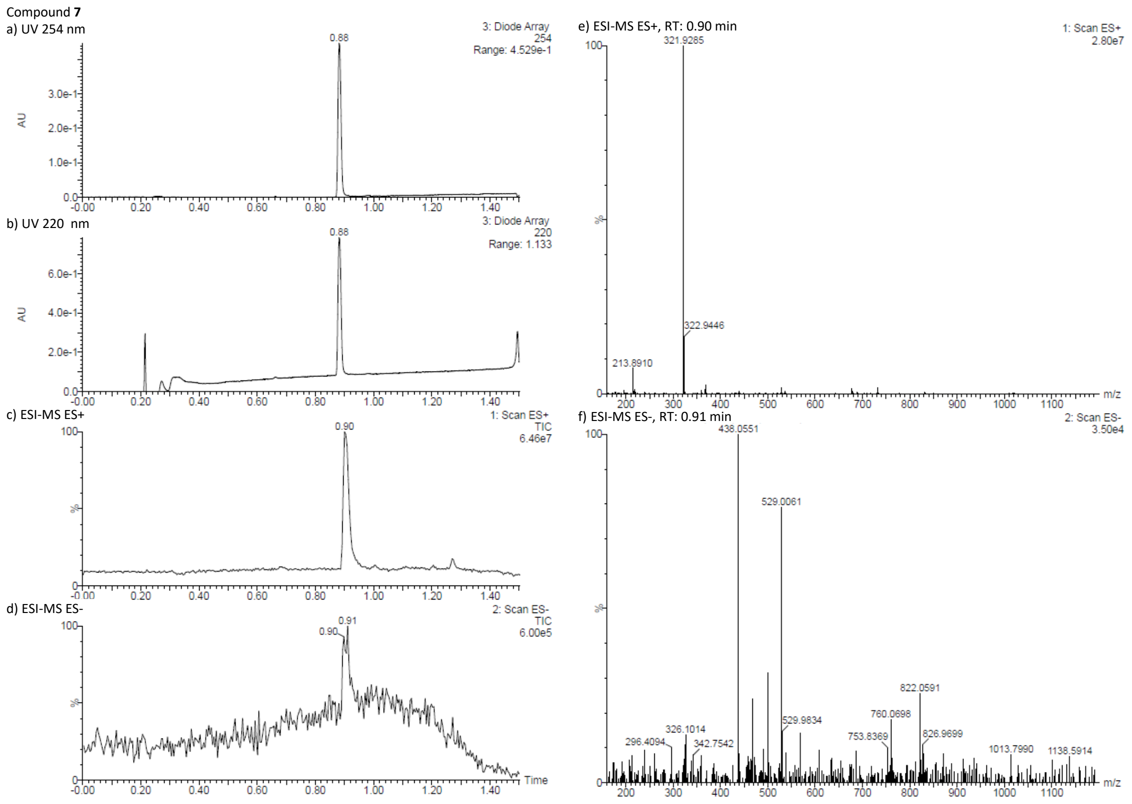

Figure S3. UPLC chromatograms of compound **7**. a) Detection by absorbance at 254 nm, b) Detection by absorbance at 220 nm, c) Scan by ESI-MS ES+, d) Scan by ESI-MS ES-, e) ESI-MS ES+ at 0.90 min, f) ESI-MS ES- at 0.91 min

### 4.4. Synthesis of amine (8)

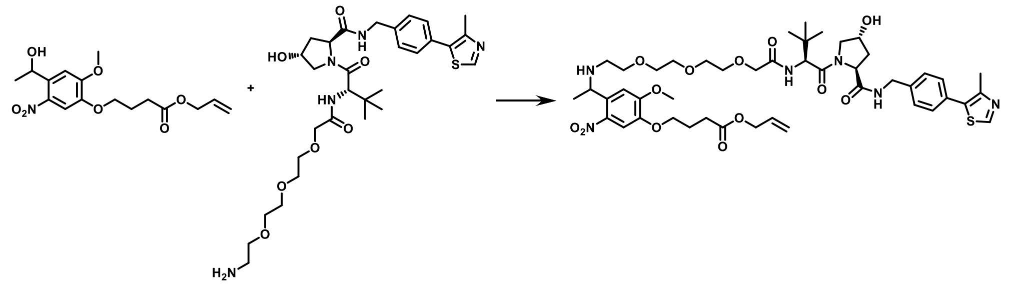

To a solution of allyl 4-[4-(1-hydroxyethyl)-2-methoxy-5-nitro-phenoxy]butanoate (**7**, 113 mg, 0.33 mmol, 1.10 equiv.) in $CH_2Cl_2$ (1.3 mL) was added $PBr_3$ (38 μL, 0.40 mmol, 1.35 equiv.) at 0 °C. The reaction mixture was stirred at rt for 15 min (pale yellow solution, SM remained, intermediate bromide observed, 1.10 min, 401.8/403.8 [M+H]+). To the mixture was added ice-water, then the mixture was diluted with EtOAc. The whole mixture was washed successively with sat.$NaHCO_3$ and brine, dried over $Na_2SO_4$, filtered and the filtrate was concentrated under reduced pressure to give a yellow residue.

(2S,4R)-1-[(2S)-2-[[2-[2-[2-(2-aminoethoxy)ethoxy]ethoxy]acetyl]amino]-3,3-dimethyl-butanoyl]-4-hydroxy-N-[[4-(4-methylthiazol-5-yl)phenyl]methyl]pyrrolidine-2-carboxamide hydrochloride (197 mg, 0.30 mmol, 1.00 equiv.) was dissolved in DMF (0.5 mL). To the solution was added $Et_3N$ (42 µL, 0.30 mmol, 1.00 equiv.). To the mixture was added a solution of intermediate bromide in DMF (1.5 mL), the reaction mixture was shaken at 60 °C for 30 min. To the mixture was added $Et_3N$ (84 µL, 0.604 mmol, 2.00 equiv.), the reaction mixture was shaken at 60 °C for 30 min. The reaction mixture was shaken at 100 °C for 8 h. The reaction mixture was concentrated under reduced pressure. HPLC preparation were performed on a Xbridge C18 (Φ30mm x 50mm, 5µm) column at a flow rate of 50 ml/min with gradient: 50% B (0 to 2 minutes), 50% to 80% (2 to 5 minutes) (buffer A = 0.05%TFA in mQ millipore water, B = 0.05%TFA in $CH_3CN$) to afford amine (**8**) (47.1 mg, 0.045 mmol, 14.9% yield) as a brown gum. To next reaction without further purifications. Amine (**8**): LC retention time: 0.75 min, MS characterization (*m/z*, $C_{46}H_{64}N_6O_{13}S$, ESI): calculated [M+H]+: 941.4325, found: 941.1465.

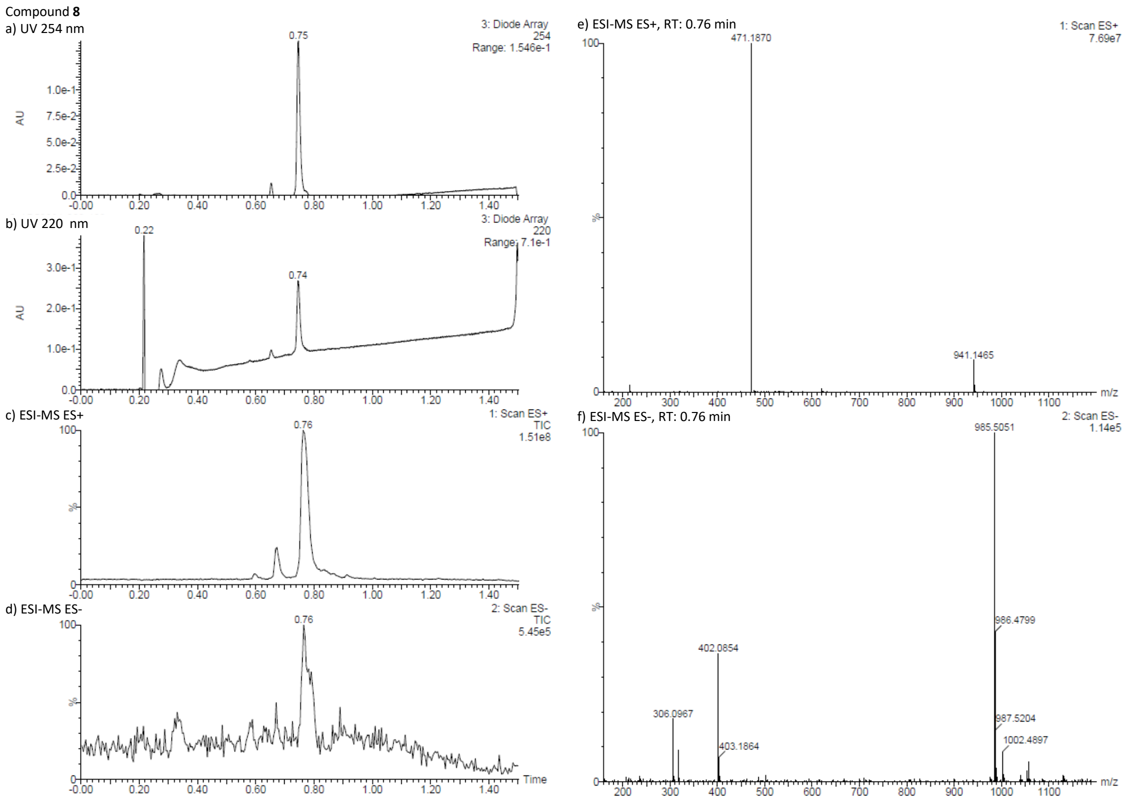

Figure S4. UPLC chromatograms of compound **8**. a) Detection by absorbance at 254 nm, b) Detection by absorbance at 220 nm, c) Scan by ESI-MS ES+, d) Scan by ESI-MS ES-, e) ESI-MS ES+ at 0.76

min, f) ESI-MS ES- at 0.76 min

### 4.5. Synthesis of PROTAC FKBP degrader-3 photo-linker allyl ester (9)

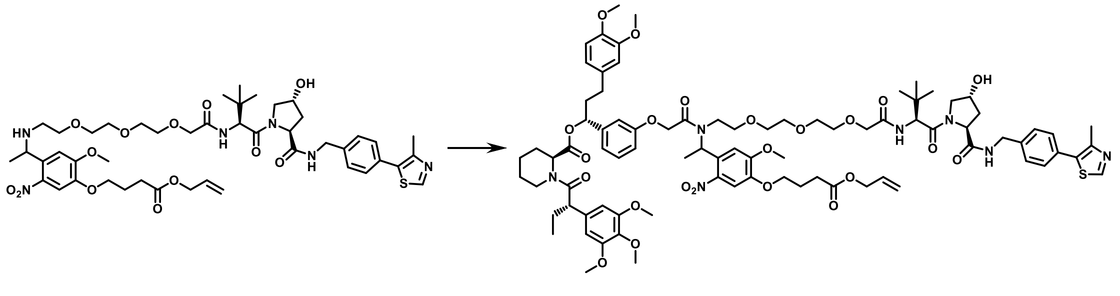

To a solution of amine (**8**) (47.1 mg, 0.045 mmol, 1.00 equiv.) and 2-[3-[(1R)-3-(3,4-dimethoxyphenyl)-1-[(2S)-1-[(2S)-2-(3,4,5-trimethoxyphenyl)butanoyl]piperidine-2-carbonyl]oxy-propyl]phenoxy]acetic acid (38 mg, 0.055 mmol, 1.23 equiv.) and HATU (21 mg, 0.055 mmol, 1.24 equiv.) in DMF (0.6 mL) was added DIPEA (19 µL, 0.11 mmol, 2.46 equiv.) at rt. The whole mixture was stirred at rt for 3 h. To the mixture was added DIPEA (19 µL, 0.11 mmol, 2.46 equiv.) and the whole mixture was stirred at 60 °C for 3 h. To the mixture was added HATU (42 mg, 0.11 mmol, 2.48 equiv.) and DIEA (40 µL, 0.23 mmol, 5.18 equiv.). The whole mixture was stirred at 80 °C for 24 hours. The mixture was diluted with EtOAc, washed with sat.$NaHCO_3$, dried over $Na_2SO_4$, filtered and the filtrate was concentrated under reduced pressure to give a residue. The residue was dissolved in DMF (0.6 mL). To the solution was added 2-[3-[(1R)-3-(3,4-dimethoxyphenyl)-1-[(2S)-1-[(2S)-2-(3,4,5-trimethoxyphenyl)butanoyl]piperidine-2-carbonyl]oxy-propyl]phenoxy]acetic acid (110 mg, 0.16 mmol, 3.55 equiv.), HATU (63 mg, 0.166 mmol, 3.71 equiv.) and DIPEA (120 µL, 0.69 mmol, 15.5 equiv.) at room temperature. The whole reaction mixture was shaken at 100 °C overnight. The reaction mixture was diluted with EtOAc, washed succesively with sat.$NaHCO_3$aq and brine, dried over $Na_2SO_4$, filtered and the filtrate was concentrated under reduced pressure to give a crude residue. HPLC preparation were performed on a Xbridge C18 (Φ30mm x 50mm, 5µm) column at a flow rate of 50 ml/min with gradient: 50% B (0 to 2 minutes), 50% to 80% (2 to 5 minutes) (buffer A = 0.05%TFA in mQ millipore water, B = 0.05%TFA in $CH_3CN$) to afford PROTAC FKBP Degrader-3 Photo-Linker allyl ester (**9**) (9.8 mg, 0.0061 mmol, 14% yield) as a brown gum. To next reaction without further purifications. PROTAC FKBP Degrader-3 Photo-Linker allyl ester (**9**): LC retention time: 1.12 min, MS characterization (*m/z*, $C_{84}H_{109}N_7O_{23}S$, ESI): calculated [M+2H]/2+: 808.8721, found: 809.0246.

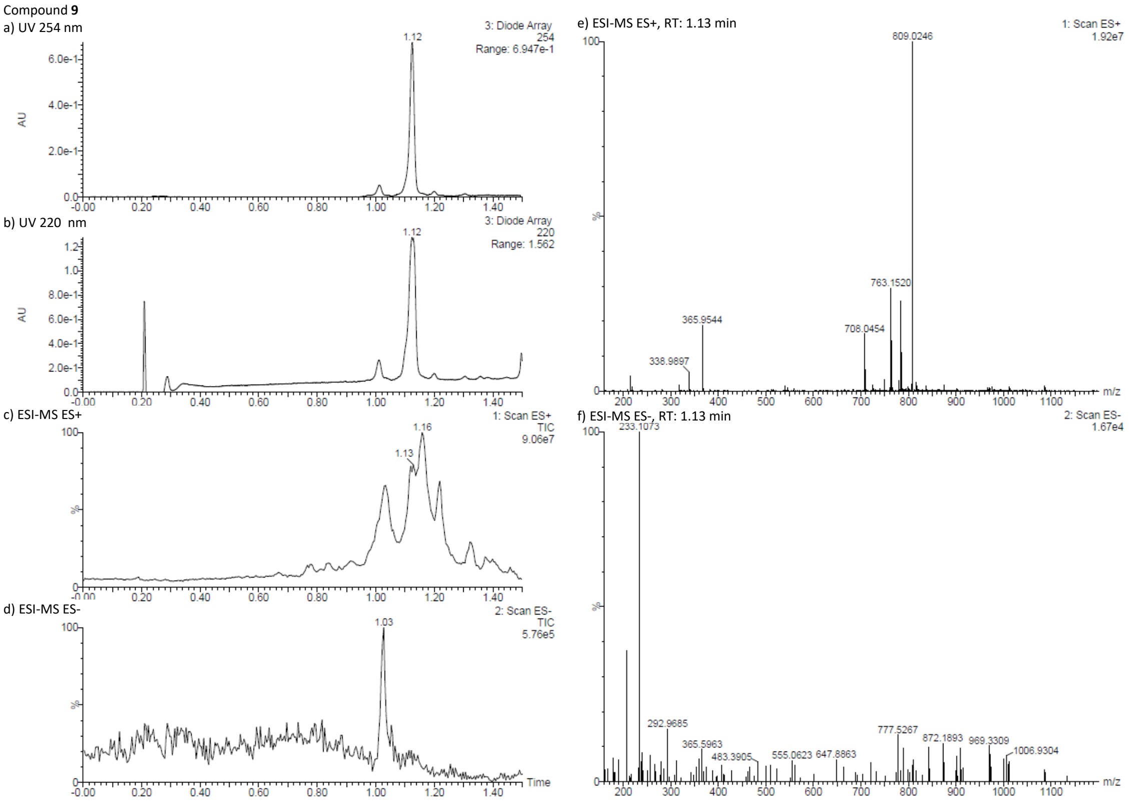

Figure S5. UPLC chromatograms of compound **9**. a) Detection by absorbance at 254 nm, b) Detection by absorbance at 220 nm, c) Scan by ESI-MS ES+, d) Scan by ESI-MS ES-, e) ESI-MS ES+ at 1.13 min, f) ESI-MS ES- at 1.13 min

### 4.6. Synthesis of PROTAC FKBP degrader-3 photo-linker carboxylic acid (10)

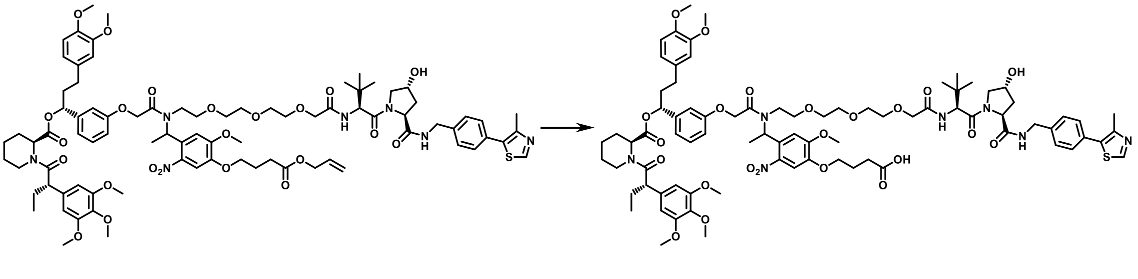

To a solution of PROTAC FKBP Degrader-3 Photo-Linker allyl ester (**9**) (9.8 mg, 0.0061 mmol, 1.00 equiv.) in $CH_2Cl_2$ (0.5 mL) was added $Pd(PPh_3)_4$ (2 mg, 0.0017 mmol, 0.28 equiv.) and 1,3-Dimethylbarbituric acid (8 mg, 0.051 mmol, 8.36 equiv.) at room temperature. The reaction mixture was shaken at room temperature overnight. The reaction mixture was concentrated under reduced pressure. HPLC preparation were performed on a Xbridge C18 (Φ30mm x 50mm, 5μm) column at a flow rate of 50 ml/min with gradient: 50% B (0 to 2 minutes), 50% to 80% (2 to 5 minutes) (buffer A = 0.05%TFA in mQ millipore water, B = 0.05%TFA in $CH_3CN$) to afford PROTAC FKBP Degrader-3 Photo-Linker carboxylic acid (**10**) (4.1 mg, 0.0026 mmol, 43% yield) as an orange gum. To next reaction without further purifications. PROTAC FKBP Degrader-3 Photo-Linker carboxylic acid (**10**): LC retention time: 1.03 min, MS characterization (*m/z*, $C_{81}H_{105}N_7O_{23}S$, ESI): calculated [M+H] +: 1576.7056, found: 1576.8063. $^1$H NMR (600 MHz, $CDCl_3$, diastereomer mixtures of rotamers) δ = 9.10 - 8.90 (m, 0.7 H), 7.87 - 7.77 (m, 0.5 H), 7.76 - 7.62 (m, 3 H), 7.61 - 7.21 (m, 30.8 H), 7.21 - 7.07 (m, 3 H), 7.01 - 6.89 (m, 2 H), 6.87 - 6.58 (m, 11 H), 6.50 - 6.33 (m, 3 H), 6.24 (s, 0.5 H), 6.01 - 5.70 (m, 1.5 H), 5.68 - 5.56 (m, 1 H), 5.53 - 5.40 (m, 2 H), 4.99 - 4.43 (m, 10 H), 4.42 - 4.23 (m, 1.5 H), 4.22 - 4.00 (m, 5 H), 4.00 - 3.91 (m, 5 H), 3.89 - 3.73 (m, 26 H), 3.71 - 3.26 (m, 41.5 H), 3.25 - 3.05 (m, 3.5 H), 3.03 - 2.77 (m, 3 H), 2.73 - 2.62 (m, 3.5 H), 2.59 - 2.41 (m, 9 H), 2.40 - 1.88 (m, 13 H), 1.83 - 1.35 (m, 11 H), 1.35 - 1.03 (m, 3 H), 1.01 - 0.92 (m, 10 H), 0.91 - 0.77 (m, 6 H). $^{13}$C NMR (150 MHz, $CDCl_3$, diastereomer mixtures of rotamers) δ = 12.48, 14.88, 18.74, 20.81, 20.86, 20.87, 20.9, 20.92, 23.91, 24.48, 25.18, 25.30, 26.34, 26.82, 27.22, 28.16, 28.27, 28.32, 29.95, 30.00, 31.26, 31.45, 31.58, 35.08, 35.10, 36.58, 36.61, 38.20, 38.25, 38.42, 40.39, 42.69, 43.10, 43.12, 43.13, 43.45, 43.55, 50.71, 51.09, 52.13, 52.22, 52.33, 55.85, 55.87, 55.95, 56.29, 56.48, 57.02, 58.95, 60.78, 60.89, 65.58, 67.11, 68.32, 69.67, 69.74, 69.80, 69.83, 69.94, 69.96, 70.07, 70.15, 70.25, 70.31, 70.48, 70.53, 70.55, 75.82, 75.88, 75.93, 75.96, 76.12, 76.29, 76.32, 76.52, 77.64, 77.68, 77.71, 77.76, 81.59, 85.94, 104.58, 104.82, 104.95, 105.01, 108.74, 110.46, 111.32, 111.34, 111.65, 111.76, 113.18, 113.58, 115.77, 119.42, 120.16, 120.22, 126.69, 127.08, 128.05, 128.13, 128.22, 128.41, 128.45, 128.48, 128.52, 128.59, 128.62, 128.67, 128.76, 128.86, 128.91, 129.15, 129.37, 129.51, 129.57, 129.65, 129.69, 130.86, 131.06, 131.31, 131.37, 131.38, 131.67, 132.14, 132.21, 132.47, 133.32, 133.54, 133.57, 134.04, 134.24, 134.31, 134.42, 134.46, 134.49, 134.54, 134.62, 134.71, 134.76, 134.80, 135.14, 135.16, 135.21, 136.51, 136.58, 142.08, 142.65, 147.31, 147.42, 147.45, 148.86, 148.91, 148.93, 148.98, 151.66, 151.68, 153.15, 153.48, 153.53, 153.62, 158.00, 158.11, 169.57, 170.05, 170.55, 170.59, 171.29, 171.42, 171.46, 173.06, 173.69.

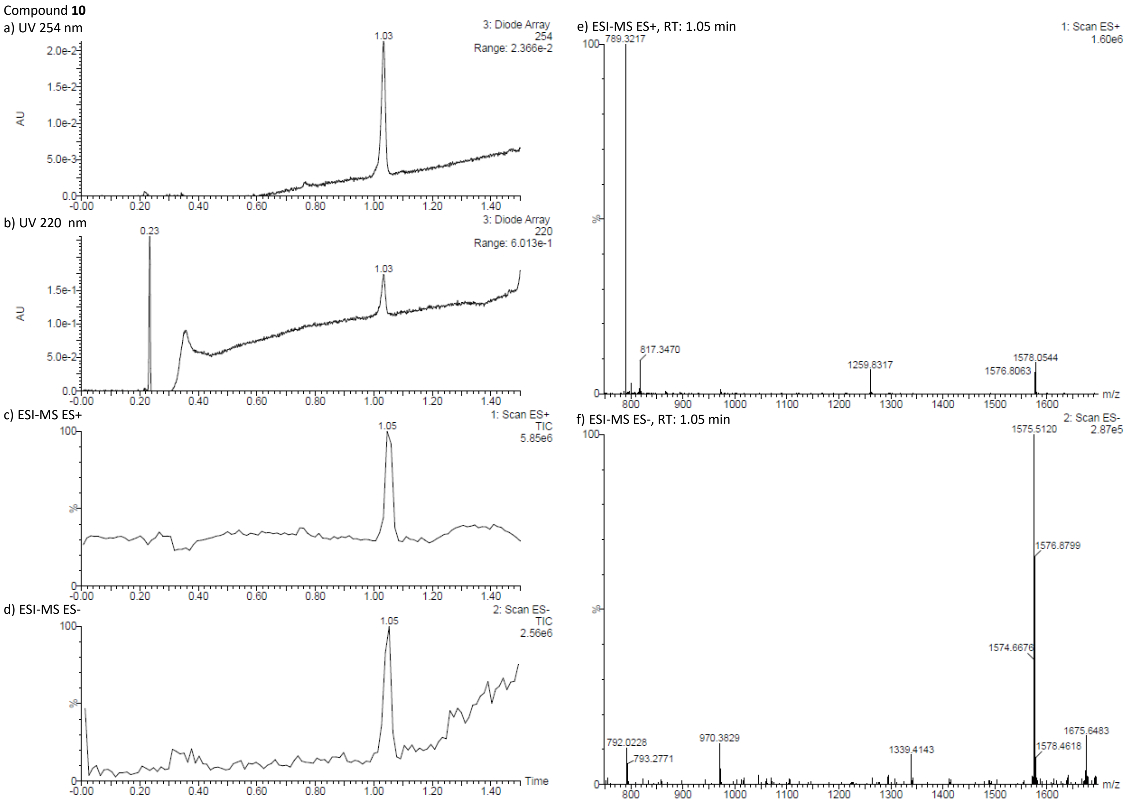

Figure S6. UPLC chromatograms of compound **10**. a) Detection by absorbance at 254 nm, b) Detection by absorbance at 220 nm, c) Scan by ESI-MS ES+, d) Scan by ESI-MS ES-, e) ESI-MS ES+ at 1.05 min, f) ESI-MS ES- at 1.05 min.

a) $^{1}H$ NMR (600MHz, $CDCl_3$)

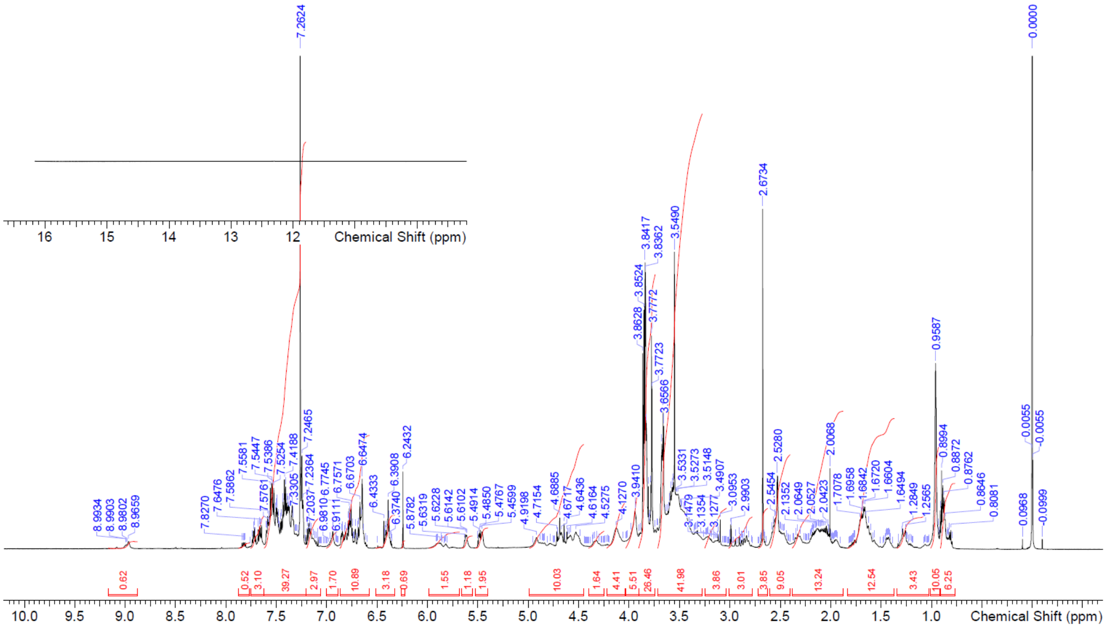

b) $^{13}C$ NMR (150MHz, $CDCl_3$)

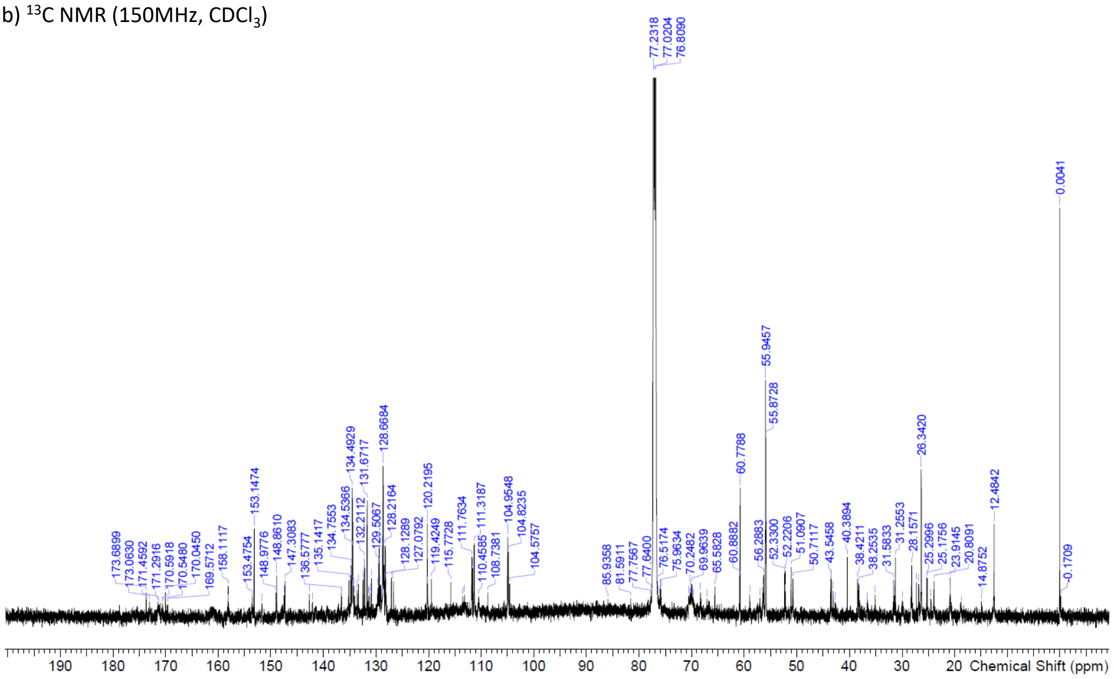

Figure S7. NMR spectra of compound **10**. a) $^{1}H$ NMR (600 MHz, $CDCl_3$, diastereomer mixtures of rotamers), b) $^{13}C$ NMR (150 MHz, $CDCl_3$, diastereomer mixtures of rotamers)

### 4.7. Synthesis of linker-modified and DNA-conjugated beads

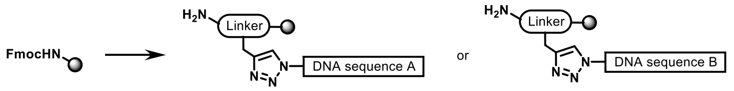

DNA-conjugated beads were synthesized from TentaGel® M RAM resin (91 mg, 20 µmol, 0.22 mmol/g) via solid-phase peptide synthesis (SPPS) followed by DNA conjugation, as previously reported by Kodadek and Paegel[2]. This procedure yielded two types of linker-modified and DNA-conjugated beads: one with DNA sequence A (**11**), and another with DNA sequence B (**6**), respectively.

### 4.8. Synthesis of PROTAC FKBP degrader-3 photo-linker OBOC-DEL (5)

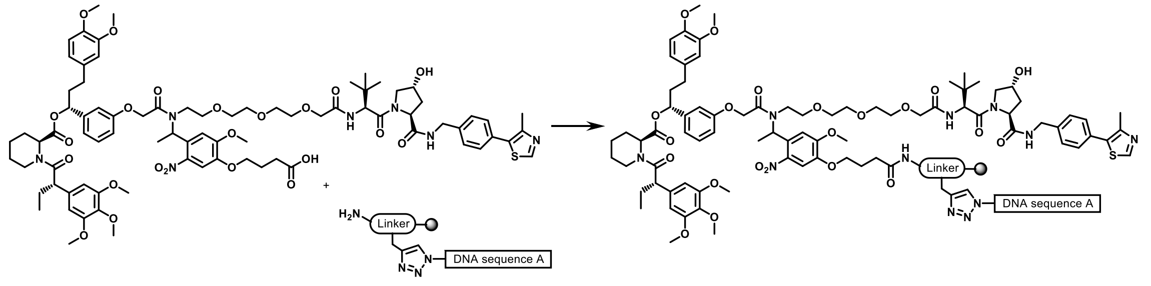

Linker-modified beads conjugated with DNA sequence A (**11**) (5 mg, 1.1 µmol, 0.22 mmol/g) was swollen with DMF (1.0 mL) in a 1 mL Mobicol "Classic" Column (M1002, MoBiTec GmbH) with inserted small filter (M2110, 10 µm pore size, MoBiTec GmbH). After 1 h, the resin was washed with DMF (×5). The resin was swollen with DMF (1.0 mL) for 10 min. FKBP degrader-3 DMF solution (4.1 mg, 2.6 µmol, 2.4 equiv., dissolved in 100 µL DMF), 0.3M DIPEA (1.4 mg, 10.8 µmol, 9.8 equiv. dissolved in 36µL NMP) and 0.3M HATU (2.3 mg, 6.0 µmol, 5.5 equiv., dissolved in 20µL DMF) were added and the mixture was shaken at room temperature. After 30 min, the resin was washed with DMF (×5) and THF (x5) to afford PROTAC FKBP Degrader-3 Photo-Linker OBOC-DEL (**5**) (5 mg, 1.1 µmol, quant.).

### 4.9. Photo-cleavage reaction of PROTAC FKBP Degrader-3 Photo-Linker OBOC-DEL (5) to yield PROTAC FKBP degrader-3 (4)

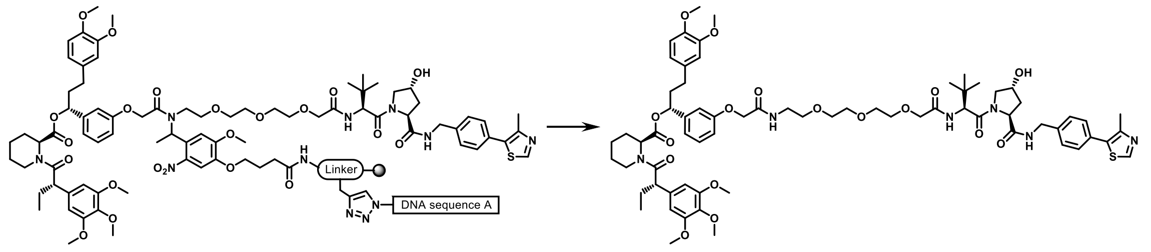

PROTAC FKBP Degrader-3 Photo-Linker OBOC-DEL (**5**) (0.1 mg, 0.022 mmol) was swollen with DMF (50 µL) in a 1.5 mL eppendorf tube. The eppendorf tube was irradiated in a well-ventilated photochemical reactor (EvoluChem PhotoRedOx Box™ and 365 nm EvoluChem™ LED, HepatoChem, MA 01915 USA) at 365 nm at room temperature. After 10 minutes, the resin was filtered, and the filtrate was analyzed via UPLCMS to detect PROTAC FKBP degrader-3 (**4**). PROTAC FKBP degrader-3 (**4**): LC retention time: 1.01 min, MS characterization (*m/z*, $C_{68}H_{90}N_6O_{17}S$, ESI): calculated [M+H]+: 1295.6156, found: 1295.7567.

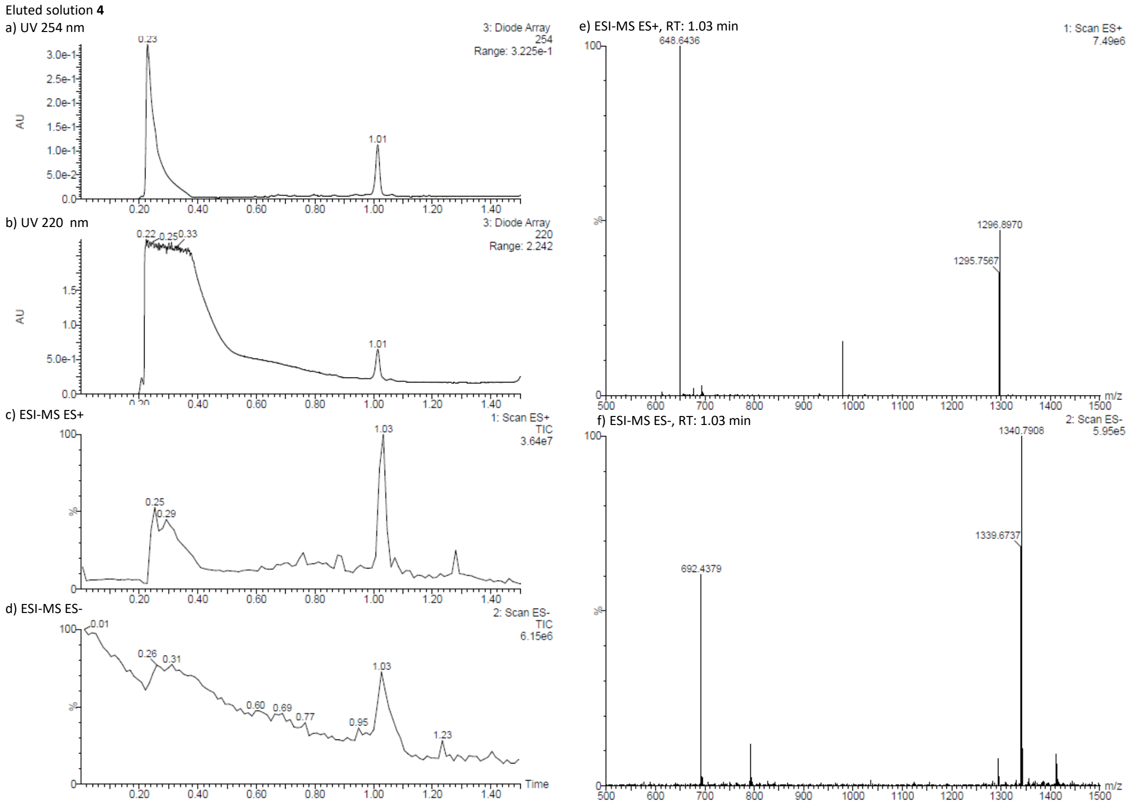

Figure S8. UPLC chromatograms of eluted solution **4**. a) Detection by absorbance at 254 nm, b) Detection by absorbance at 220 nm, c) Scan by ESI-MS ES+, d) Scan by ESI-MS ES-, e) ESI-MS ES+ at 1.03 min, f) ESI-MS ES- at 1.03 min

### 4.10. Structure of PROTAC FKBP degrader-3 photo-linker OBOC-DEL (5) (positive control) and compound unconjugated OBOC-DEL (6) (negative control)

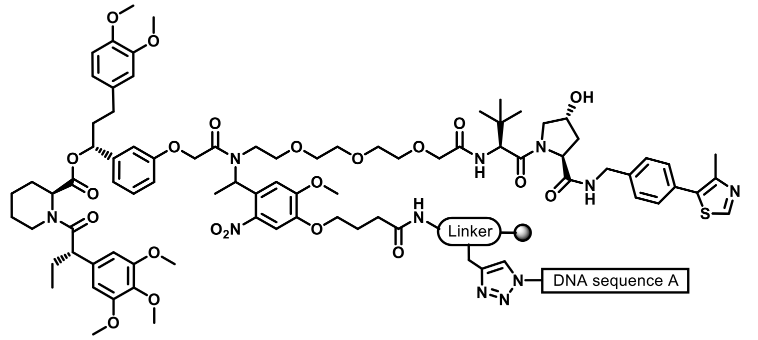

PROTAC FKBP degrader-3 photo-linker OBOC-DEL (**5**)
Positive control

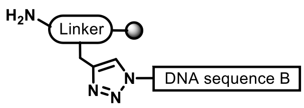

Compound unconjugated OBOC-DEL (**6**)
Negative control

### 5. Study of compound separation from beads in the Beacon system

#### 5.1. Optimization of bead load conditions

After several examinations in actual experiments, the conditions for loading OBOC-DEL beads into the Beacon were determined as follows:

Bead concentration was adjusted to $1.5 \times 10^6$ particles/mL in PBST and dispersed by pipetting thoroughly just before Load. Parameters for Load were changed from the default values to 25µL for Import Volume, 3 times for Pen Chip, and Target and Pen Selection (TPS) for Penning Algorithm. TPS was set for each experiment so that particles were transported to the selected the Field of View (FOV) and the Pens. The OEP parameters were set to 10.0um/s, 7.0V, and 1.3MHz for Cage Speed, Waveform Voltage, and Waveform Frequency, respectively. Beads were loaded at a temperature of 25 deg.C. Since it was found that the size of the beads varied a little depending on the type of bead synthesis (production lot), the optimal Cell Counting Algorithm was selected according to the bead. For the synthetic beads used in this study, CNN Cell Detect was applied. By changing the Cell Counting Algorithm, the bead penning efficiency and counting accuracy were dramatically improved.

#### 5.2. Time settings for time-lapse imaging

To estimate the appropriate imaging timing when using the Beacon system, the Wilke–Chang equation—commonly employed in chemical engineering—was applied. Calculations were performed assuming an incubation temperature of 36°C. The height of the nanopen on the 3500 chip is 370 µm, and the diffusion rate was calculated over this distance. Assuming the same substance is present in both the channel and the nanopen, the flow rate in the channel was not considered. The molecular weight was assumed to be 400 Da, based on the average molecular weight of compounds used in Activity-Based DNA Encoding Library Screening[3]. While the Wilke–Chang equation is useful for estimating diffusion coefficients of unknown substances, it is not universally applicable and should be used as a reference.

Wilke-Chang equation: $D \cdot \eta / T = 4.98 \times 10^{-14} M_{w}^{-0.6}$

$\eta = 7 \times 10^{-4}$ [Pa·s] (viscosity of water at 36°C)

$T = 273+36$

The diffusion coefficient $D$ is estimated to be 4.23E-10[$m^2$/s]. Based on this, the time required for diffusion from the 370 µm nanopen to the channel is approximately 2 minutes.

Therefore, imaging was initiated at least 2 minutes after UV irradiation. In practice, the imaging interval was adjusted for each experiment, ranging from every 30 minutes to every 2 hours. For cell phenotype evaluation, time-lapse imaging was conducted for up to 6 hours, referencing the time point at which the effects of the target compound on the cells were observed in prior manual experiments.

### 5.3 Adjustment of FITC exposure time and creating calibration curves

Carboxyfluorescein solutions were prepared at the desired concentrations and used for fluorescence intensity measurements. Fluorescence intensity in the FITC channel was acquired at various exposure times to investigate the relationship between concentration and exposure duration (Figure S9A). Based on these results, a calibration curve was constructed to estimate approximate fluorescein amide concentrations from fluorescence intensity at specific exposure times (Figure S9B). Furthermore, equations were derived from the calibration curves to calculate concentrations for each exposure time, which were used in subsequent experiments (Figure S9C).

Assuming complete cleavage of all linkers from OBOC-DEL beads within a nanopen, the estimated concentration exceeds 100 μM.

Standard capacity /bead: $1.3 \times 10^{-4}$ nmol/bead

Nanopen volume: 0.76μL

$1.3 \times 10^{-4} / 0.76 \approx 1.7 \times 10^{-4} = 170$ μM

However, excessively high drug concentrations may induce cytotoxicity. Therefore, screening is typically conducted at lower concentrations to minimize such risks. Complete linker cleavage is not essential; even a concentration of 10 μM—approximately 10% of the estimated maximum—is sufficient to induce phenotypic changes in cells.

In this screening, conditions that allow visualization at optimal brightness up to 50 μM are desirable. Our observations indicate that exposure times exceeding 750 milliseconds (ms) may result in overexposure at concentrations above 50 μM. Accordingly, to ensure accurate and reliable imaging, the FITC exposure time was set to 500 ms.

**A**

| Reagent Concentration | FITC Exposure Time (msec) | | | | | |
|---|---|---|---|---|---|---|
| | 50 | 100 | 250 | 500 | 750 | 1000 |
| 0.1 uM | 67.4 | 148 | 354 | 745.1 | 1150.9 | 1475.6 |
| 1.0 uM | 144.5 | 276.1 | 713.7 | 1445.7 | 2167.9 | 2762.0 |
| 10.0 uM | 739.7 | 1507.4 | 3898.8 | 7853.3 | 11927.6 | 16182.3 |
| 50.0 uM | 4523.9 | 8940.3 | 22435.5 | 44418.7 | 62582.2 | 62989.4 |
| 100.0 uM | 10545.7 | 20916.6 | 52844.2 | 62966.5 | 62983.3 | 62883.7 |

**B**

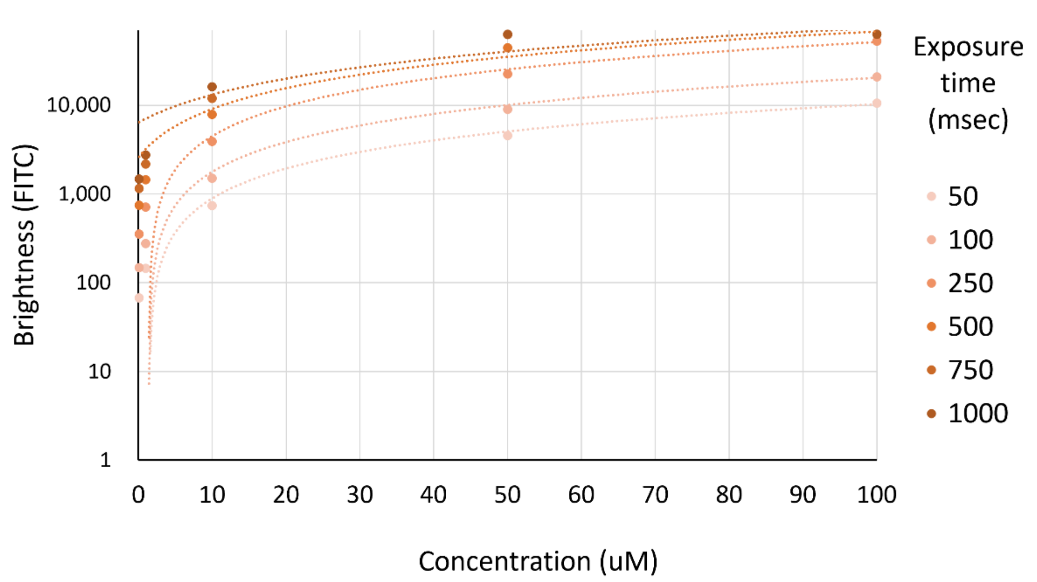

**C**

| FITC Exposure Time | Concentration Calculation Formula | $R^2$ |
|---|---|---|
| 50 msec | y = 104.19x - 152.87 | 0.9942 |
| 100 msec | y = 206.44x - 293.91 | 0.9941 |
| 250 msec | y = 520.95x - 735.63 | 0.9937 |
| 500 msec | y = 649.64x + 2554.5 | 0.9640 |
| 750 msec | y = 676.36x + 6370.2 | 0.8337 |
| 1,000 msec | y = 659.75x + 8001.4 | 0.8242 |

Figure S9. (A) Fluorescein fluorescence intensity values measured at various exposure times during Fluorescein fluorescence imaging for each reagent concentration. The data suggests that signal saturation may occur at exposure times exceeding 750 ms for 50 µM and 500 ms for 100 µM fluorescein. (B) Calibration curves constructed based on the results in (A), illustrating the relationship between fluorescein concentration and fluorescein fluorescence intensity for each exposure time. Each curve corresponds to a different exposure duration (50 - 1,000 ms), enabling quantitative estimation of fluorescein concentration from measured brightness values. (C) Calibration equations derived from the curves in (B) for each exposure time. Each equation represents a linear relationship between fluorescence intensity (x) and fluorescein concentration (y), with the corresponding $R^2$ values indicating the goodness of fit. In subsequent experiments for linker cleavage validation, the equation for 500 ms exposure was applied to estimate the concentration of the released compound.

For the bead loading and imaging experiments on the microfluidic chip, a series of sequential operations were integrated to establish the following workflow (see Table S1).

Table S1. Beacon Workflow and Imaging Settings for Confirming Linker Cleavage in OBOC-DEL Beads

| Process | Operations | Details |
|---|---|---|
| Preparation | Prime Line and Chip w/Media | The fluid inside the chip containing the channels and nanopens was replaced with PBST solution which was used to condition the beads. |
| Beads Load | Load | Import beads to chip and pen a single bead to nanopen.<br>Pen Chip <3 times - Unpen Multiple |
| $CO_2$ Blow | Manual operation | Inject $CO_2$ into the channel of the chip and remove as much liquid as possible from the channel. |
| Linker photoinduced dissociation & Imaging | Multi-Spectral Time Lapse Imaging | One FOV of the chip was illuminated with UV (390 nm)* and the fluorescence attached to the cleaved linkers was detected with FITC.<br>*DAPI channel was used instead of UV. |
| | TPS Imaging | Chip calibration - Imaging (OEP, FITC 500 ms) - UV Exposure (10 sec) - Imaging - UV Exposure - Imaging -... Repeat within a specified time. |
| Cleaning | Flush<br>Culture | Flow the PBST solution or culture medium through the chip again. After washing out any remaining particles in the channel at the maximum flow rate, continue to flow the liquid in small amounts to prevent the chip from drying out. |
| Analysis | - | Analyze the acquired images using Image Analyzer to calculate bead counts and brightness. |

In our previous experiments, we confirmed that 10 minutes of UV irradiation can cleave the linker from OBOC-DEL beads. Since the near-UV light (UV-A) transmitted through the DAPI filter (excitation wavelength: 390/40 nm) installed in the Beacon system is the closest alternative to direct UV irradiation, we used this light source instead. However, the Beacon system allows irradiation for a maximum of 10 seconds per cycle. Therefore, to ensure sufficient linker cleavage, we repeated the irradiation with 390 nm light multiple times (Figure S10).

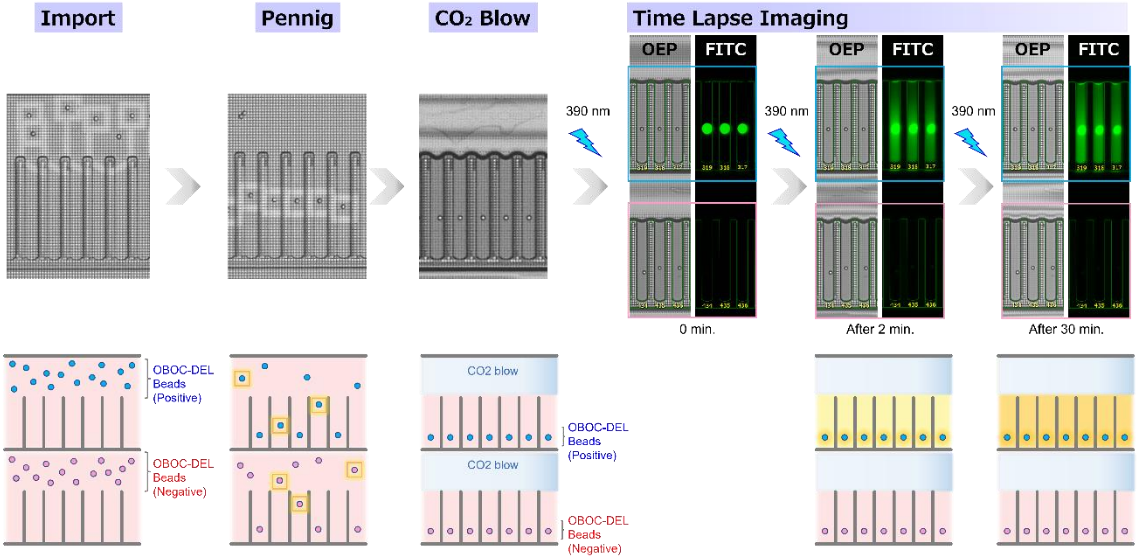

Figure S10. Workflow of the experiment using the Beacon system: OBOC-DEL beads adjusted to a concentration of 1.5×10⁶ particles/mL were loaded into the OptoSelect™ chip and individually positioned into nanopens using Opto-ElectroPositioning (OEP) technology. Residual liquid in the channel was removed manually using $CO_2$ flow. Linker cleavage was induced by irradiating with light passed through a DAPI filter (excitation wavelength: 390 nm) for 10 seconds per cycle. fluorescein fluorescence from the released compounds was monitored at defined intervals using time-lapse imaging, with each imaging session preceded by 10 seconds of irradiation.

### 5.4 Comparison of positive and negative beads

To evaluate the extent of linker dissociation induced by UV (390 nm) irradiation, fluorescein - labeled linker-conjugated beads (positive beads) and unconjugated beads (negative beads) were introduced into separate nanopens within the same field of view (FOV). Fluorescein fluorescence was observed from the positive beads within one minute, and the intensity gradually increased over time. After 10 minutes, the Fluorescein fluorescence stabilized at approximately 15 µM, which exceeds the threshold concentration of 10 µM required for cellular evaluation, indicating that the target concentration was sufficiently achieved. In contrast, no changes in fluorescein fluorescence intensity were observed over time in the negative beads (Figure S11). These results demonstrate the feasibility of using OBOC-DEL on the Beacon system for validation. Further optimization of experimental conditions was pursued to enable cell-based evaluation.

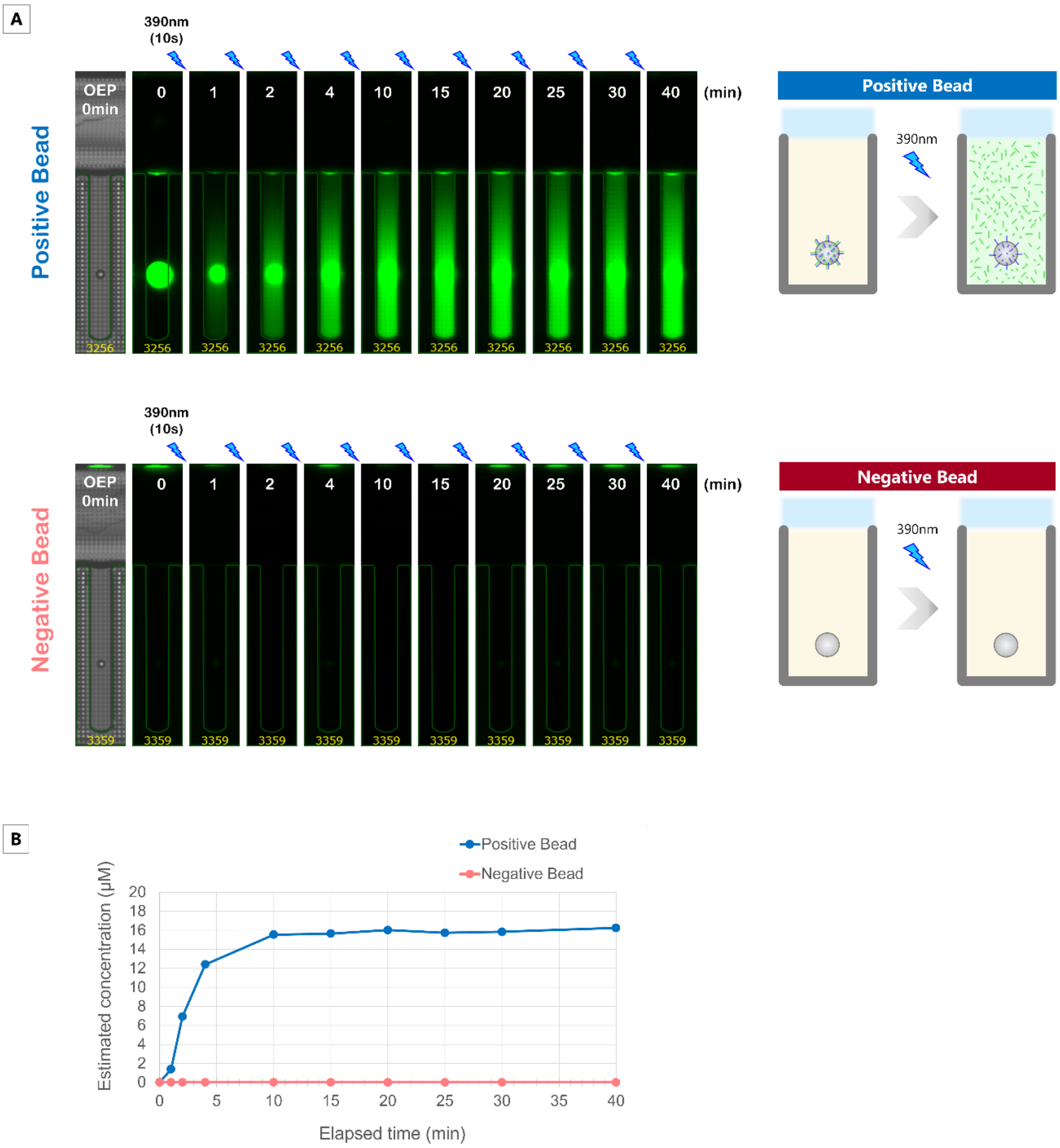

Figure S11. Fluorescein fluorescence Intensity Changes in Positive and Negative Beads. (A) Top: Positive beads conjugated with fluorescein -labeled linkers; Bottom: Negative beads without any linker conjugation. (B) Upon UV (390 nm) irradiation, fluorescein fluorescence was observed in the positive beads within 1 minute, with intensity gradually increasing over time and stabilizing at a constant level after 10 minutes. No fluorescein fluorescence was observed in the negative beads throughout the experiment.

### 5.5. Optimization of UV irradiation and assay conditions

Initial linker cleavage was performed using repeated cycles of 10-second UV exposure followed by FITC imaging. However, prolonged exposure to short-wavelength light poses a risk of cellular damage, making this approach unsuitable for cell-based assays. During a compound excision experiment, an unexpected interruption occurred after the 0-minute imaging point. When resumed 20 minutes later, fluorescein fluorescence exceeded 50 μM (Figure S12A), despite only a single 200-ms irradiation through the DAPI filter prior to the interruption. This indicates that effective cleavage can be achieved with a brief 200-ms exposure, rather than the previously assumed 10-minute duration based on tube-based experiments. Given that near-UV light (390/40 nm) may harm cells, minimizing exposure is critical. Subsequent studies focused on defining the shortest exposure time required to achieve sufficient cleavage while limiting cellular impact.

This incident also highlighted the influence of $CO_2$ flow on compound retention. Previously, $CO_2$ was continuously blown during the assay to remove residual liquid from the channels; however, this likely caused cleaved compounds to diffuse downstream into adjacent nanopens, reducing local concentration within the nanopen. In a comparative test evaluating the presence or absence of $CO_2$ blowing, beads were loaded, followed by UV irradiation (200 ms) and time-lapse imaging (Figure S12B). When $CO_2$ blowing was stopped after bead loading, the fluorescein concentration reached approximately 63 μM at 60 minutes, compared to about 12 μM under continuous blowing—an increase of roughly fivefold (Figure S12D). These results confirm that halting $CO_2$ flow prevents unintended compound leakage and promotes accumulation within the nanopen.

Overall, these findings demonstrate that Beacon enables efficient cleavage with only 200 ms of UV exposure—a standard DAPI imaging setting with minimal cellular impact—while prior removal of channel liquid and cessation of $CO_2$ flow ensure compound retention in nanoliter-scale nanopens. Additional tests using 20-ms exposure revealed slower fluorescence increase (Figure S12C): ~10 μM at 30 minutes and ~80 μM at 180 minutes, compared to >90 μM at 120 minutes with 200 ms (Figure S12E). Shorter exposure allows gentler cleavage and concentration control; therefore, 200 ms was selected to achieve target levels within ~30 minutes.

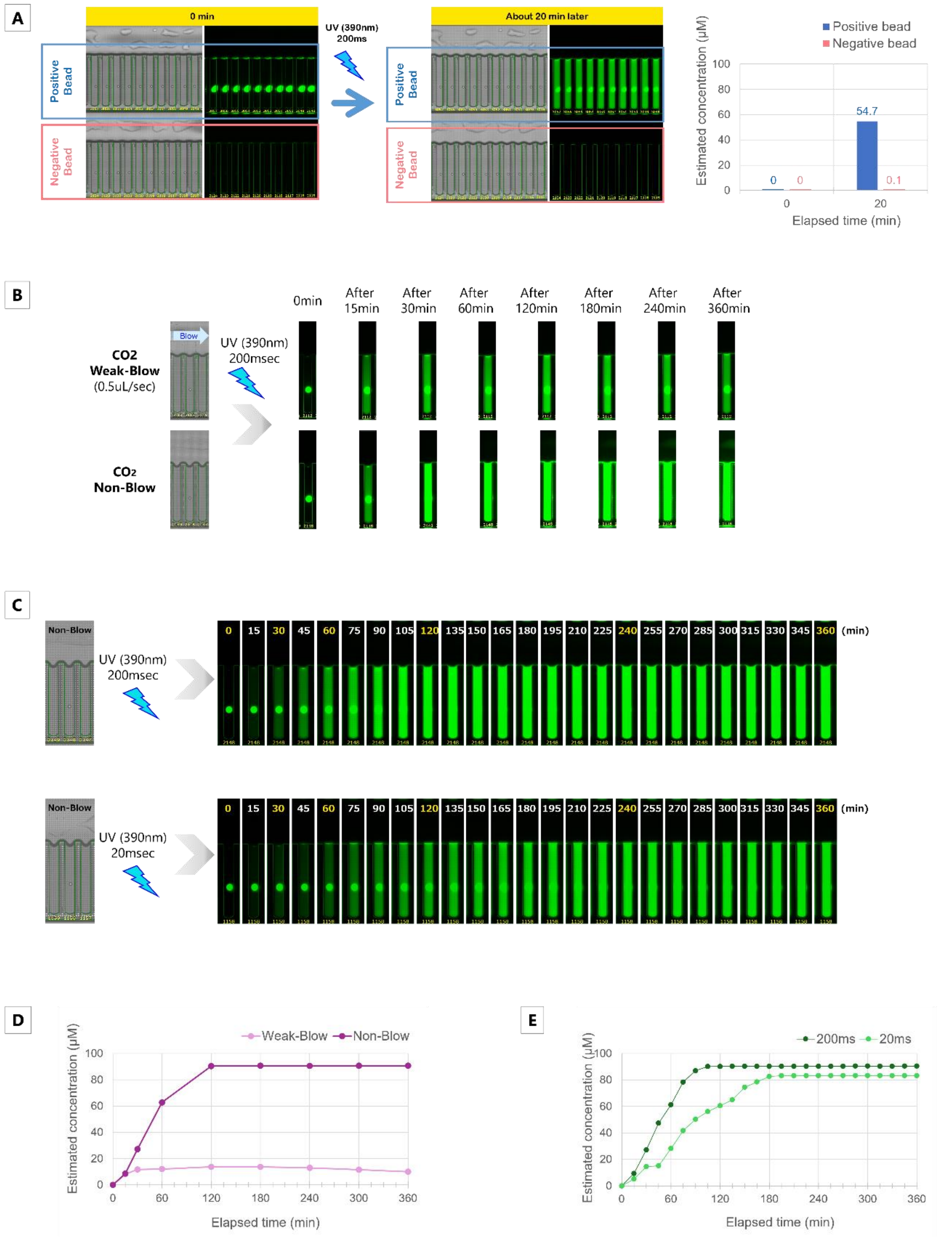

Figure S12. (A) Images acquired before and 20 minutes after a single 200-ms UV (390 nm) exposure. In the nanopen loaded with positive beads, fluorescein fluorescence indicating diffusion of the cleaved compound was observed, and the fluorescein concentration exceeded 50 μM. No changes were observed in the negative beads before and after UV exposure.

(B) Differences in the accumulation of compounds cleaved from beads with or without $CO_2$ blowing.

(C) Temporal changes in fluorescein fluorescence intensity extracted from beads under varying UV (390 nm) exposure durations. (D) Estimated fluorescein concentration over time following UV exposure, with or without $CO_2$ blowing during the assay. Halting $CO_2$ flow enabled higher accumulation of cleaved compounds within the nanopen. (E) Estimated fluorescein concentration over time following UV exposure under each irradiation condition. Compared to 200 ms, 20 ms resulted in more gradual compound cleavage.

### 5.6. Creation of bead-based workflows

Based on our previous investigations, the compound excision experiment from OBOC-DEL beads using the Beacon system was carried out according to the following procedure (see Figure S13):

1) Bead Loading into nanopens. OBOC-DEL beads were imported into the Beacon chip and individually positioned into nanopens using Opto-Electropositioning (OEP) technology.
2) Liquid Removal by $CO_2$ Blowing. To remove residual liquid from the channels, $CO_2$ was manually blown through the system for over 30 minutes. Once complete removal was confirmed, the $CO_2$ flow was stopped.
3) UV Irradiation and Fluorescence Imaging. Near-UV light (390 nm) was irradiated through a DAPI filter for 200 ms to cleave the linker and release the compound from the beads. To monitor fluorescein compounds bound to positive beads and EGFP-expressing cells, FITC exposure was set to 500 ms, and time-lapse imaging was performed for up to 6 hours.
4) Image Analysis. Captured images were imported into the Image Analyzer software, and EGFP fluorescence intensity was quantified. The intensity data were applied to a calibration curve created from known concentrations to estimate the concentration of the excised compound.
5) Bead Recovery. Target beads, including positive and negative beads, as well as those associated with cells showing EGFP fluorescence loss, were unloaded from nanopens and transferred to a 96-well plate. Barcode sequences were then amplified by PCR and identified via next-generation sequencing (NGS).

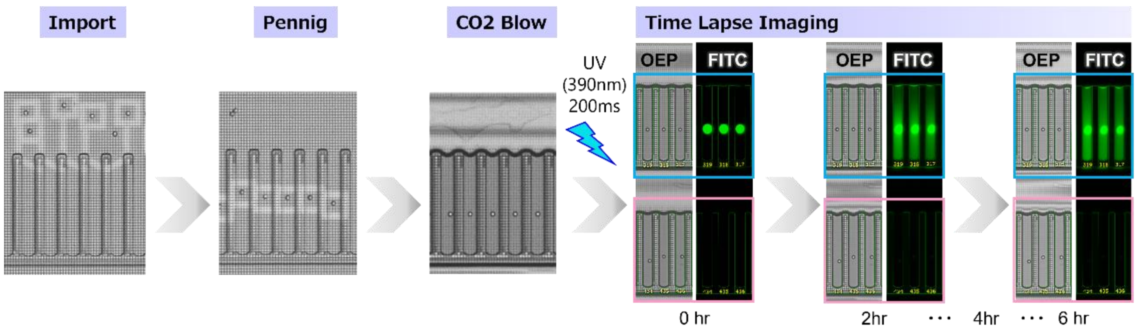

Figure S13. Optimization of the linker cleavage process using the Beacon system with OBOC-DEL beads. UV irradiation on the beads was shortened to a single 200ms pulse. The fluorescence intensity of cleaved fluorescein was extracted from images acquired at defined intervals.

To facilitate bead loading and imaging experiments on the microfluidic chip, a series of sequential operations were integrated into a standardized workflow (refer to Table S2).

Table S2. Optimized Beacon Workflow and Settings for Linker Cleavage from OBOC-DEL.

| Process | Operations | Details |
|---|---|---|
| Preparation | Prime Line and Chip w/Media | The fluid inside the chip containing the channels and nanopens was replaced with PBST solution which was used to condition the beads. |
| Beads Load | Load | Import beads to chip and pen a single bead to nanopen.<br>Pen Chip <3 times - Unpen Multiple |
| $CO_2$ Blow | Manual operation | Inject $CO_2$ into the channel of the chip and remove as much liquid as possible from the channel. |
| Linker photoinduced dissociation & Time Lapse Imaging | Multi-Spectral Time Lapse Imaging | One FOV of the chip was illuminated with UV (390 nm)* and the fluorescence attached to the cleaved linkers was detected with FITC.<br>*DAPI channel was used instead of UV. |
| | TPS Imaging | Chip calibration - Imaging (OEP, FITC 500 ms) - UV Exposure (200 ms) - Time Lapse Imaging (OEP, FITC 500 ms /6 hrs) |
| Cleaning | Flush<br>Culture | Flow the PBST solution or culture medium through the chip again. After washing out any remaining particles in the channel at the maximum flow rate, continue to flow the liquid in small amounts to prevent the chip from drying out. |
| Analysis | - | Analyze the acquired images using Image Analyzer to calculate bead counts and brightness. |
| Beads Unload | Unload | Selected beads were unpenned from the nanopen and transferred to the 96-well collection plate. |

### 6. Verification of the effect of model compounds on cellular phenotype

To evaluate whether the constructed cells could survive without issues after being loaded onto the chip, and whether EGFP expression could be observed using the Beacon system, a series of tests were conducted.

#### 6.1. Evaluation of cell loading conditions

Although PC-3 (ATCC, #CRL-1435) cells are adherent by nature, they were detached from the culture plate and handled in a spherical form to enable loading into the chip. After examining the cell loading conditions, the following procedure was established:

Cells were adjusted to a concentration of $1.5\times10^6$ cells/mL in culture medium and passed through a 40 μm cell strainer immediately before loading. They were then gently dispersed by pipetting. The loading parameters were modified from the default settings: Import Volume was set to 25 μL, Pen Chip was repeated three times, and the Penning Algorithm used was Target and Pen Selection (TPS). TPS was configured in each experiment to ensure that cells were transported to the selected Field of View (FOV) and pens. The OEP parameters were set to a Cage Speed of 8.0 μm/s, Waveform Voltage of 4.7 V, and Waveform Frequency of 1.3 MHz. Lowering the Cage Speed improved cell penning efficiency. Cell loading was performed at 36°C. The Cell Counting Algorithm applied was CNN Watershed.

#### 6.2. Cellular behavior on the microfluidic chip

Cells were loaded onto the chip and monitored for 16 hours without UV irradiation, using imaging with a 2,000 ms exposure in the FITC channel. As a result, cell proliferation was observed 6 hours after loading, and by 16 hours, approximately 35% of the nanopens showed evidence of cell doubling. Changes in EGFP fluorescence intensity for each cell were recorded at 2-hour intervals. At time 0, EGFP fluorescence intensity varied among cells, likely due to differences in cell cycle stages and EGFP expression levels. During the 16-hour observation period, EGFP fluorescence intensity fluctuated by up to approximately 6,000, but none of the cells dropped below 5,000 (Figure S14). These changes in EGFP fluorescence intensity are considered to be influenced, in part, by cell doubling. Based on these findings, a EGFP fluorescence intensity threshold of 5,000 was set prior to the assay, and only cells exhibiting EGFP fluorescence above this threshold were selected for evaluating the effects of compounds.

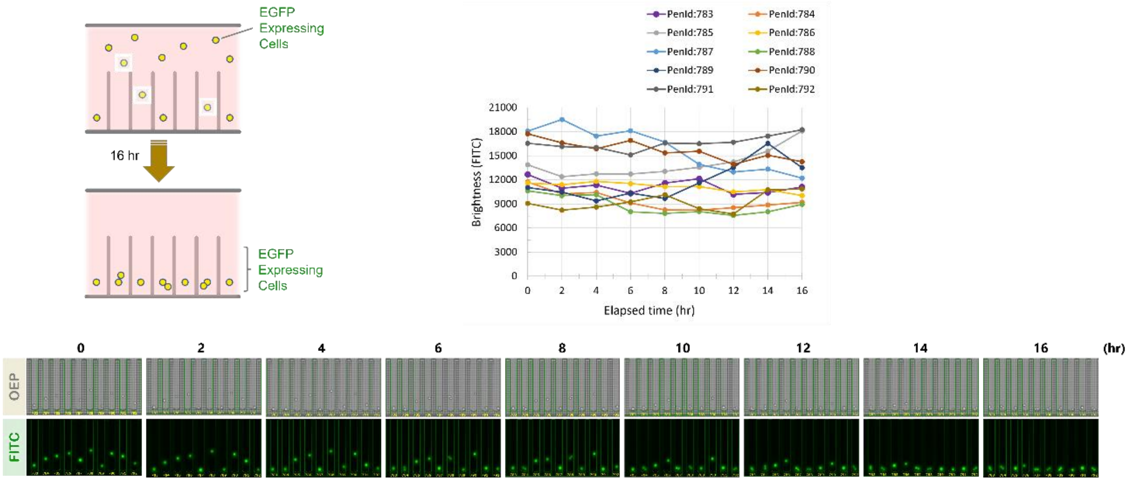

Figure S14. Single-cell culture of EGFP-expressing cells on the OptoSelect chip. Single cells were loaded into nanopens on the OptoSelect chip and monitored every 2 hours over a 16-hour period. Time-lapse imaging confirmed sustained EGFP fluorescence in most cells throughout the observation. Approximately 13% of the nanopens showed proliferating cells at 6 hours, and about 35% at 16 hours. Quantitative analysis of EGFP fluorescence intensity in individual cells showed that the average intensity remained nearly constant, with no significant decrease observed during the 16-hour observation period.

### 6.3. Observation of EGFP fluorescence disappearance in cells induced by a model compound

To confirm the disappearance of EGFP fluorescence expressed in cells, a medium containing a model compound was introduced into the chip, and the cells were cultured and observed over a 6-hour period. As a positive control compound, PROTAC FKBP Degrader-3 (MedChemExpress, #HY-135345) was added to the culture medium at a final concentration of 1.0 μM. As a result, while cell proliferation was observed over time, EGFP fluorescence intensity gradually decreased in all cells, with a marked reduction observed at 2 hours. By 6 hours, EGFP fluorescence was nearly undetectable. Prior to the assay, the initial EGFP fluorescence intensity varied widely among individual cells, but the pattern of EGFP fluorescence decline was consistent across cells (Figure S15).

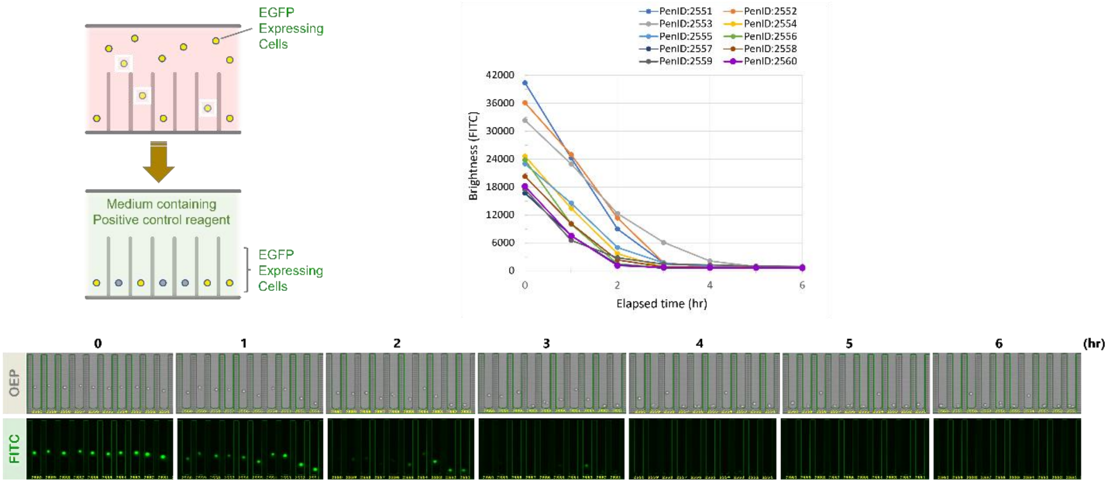

Figure S15. Time-lapse imaging and EGFP fluorescence monitoring of EGFP-expressing cells cultured on the OptoSelect chip in the presence of a positive control compound. After cell loading, the medium was replaced with one containing PROTAC FKBP Degrader-3 (1.0 μM). Cell proliferation was observed as early as 2 hours after the start of imaging. At the same time point, EGFP fluorescence intensity showed a marked decrease, and by 6 hours, EGFP fluorescence was no longer detectable in all cells.

Continuous overnight observation revealed that none of the cells exhibited significant recovery of the previously diminished EGFP fluorescence intensity. After approximately 22 hours, to assess cell viability, the culture medium was replaced with one containing SYTOX Orange Nucleic Acid Stain (Molecular Probes, #S-11368) at a final concentration of 2.0 μM, and imaging was performed using the TRED channel with a 1,000-millisecond exposure time. As a result, although cells appeared to be slightly stained under observation with the TRED channel, bright-field imaging using the OEP channel revealed clear cell contours and no morphological deformation. Furthermore, cell proliferation was observed in approximately 43% of the nanopens. These findings suggest that the cells are viable under single-cell culture conditions on the chip. (Figure S16). The PC-3 cells engineered in this study to express EGFP, although inherently adherent, demonstrated the ability to withstand one day of culture on the chip even under conditions in which EGFP expression was suppressed by the positive control compound. These findings indicate that this cell line is well-suited for compound evaluation using the Beacon system.

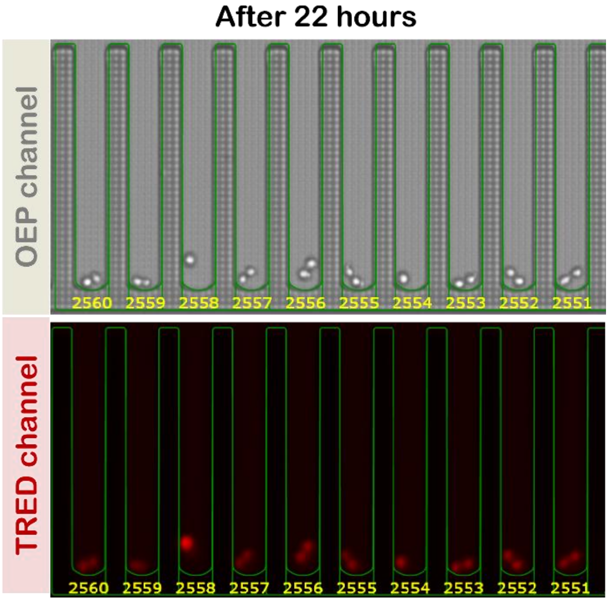

Figure S16. Assessment of cell viability after 22 hours revealed nonspecific staining by the viability assay reagent (TRED channel); however, cells exhibited clear contours and no morphological changes (OEP channel). Furthermore, cell proliferation was observed in over 43% of the nanopens, indicating that cell viability was maintained even under prolonged single-cell culture conditions on the chip.

Subsequently, experimental conditions for a small-molecule compound screening assay based on cellular phenotypic changes were optimized. A culture medium containing both the positive control compound PROTAC FKBP Degrader-3 and the viability assay reagent SYTOX Orange Nucleic Acid Stain was prepared in advance, with final concentrations of 1 µM and 2 µM, respectively. Cells were cultured for up to 6 hours, during which a sufficient decrease in EGFP fluorescence intensity was observed. Following a single 200 ms UV (390 nm) irradiation, images were acquired every 2 hours using exposure times of 2,000 ms for the FITC channel and 1,000 ms for the TRED channel. Bright-field imaging using the OEP channel revealed that approximately 12% (11-14%) of nanopens exhibited cell proliferation during the 6-hour culture period, and over 99% of cells showed no signs of damage or cell death. Cell death was confirmed in 0–1 nanopens per field of view (0–0.9%), where bright-field imaging revealed loss of cellular boundaries and morphological deformation, and TRED channel imaging showed strong fluorescence from the viability assay reagent spreading throughout the nanopen. The rate of EGFP fluorescence decay varied among individual cells, with some showing a rapid decrease in EGFP fluorescence intensity and others a more gradual decline. However, by 6 hours, strong EGFP fluorescence was no longer detected in any nanopen (Figure S17). When SYTOX Orange Nucleic Acid Stain was added to the culture medium from the beginning, saturation occurred in the TRED channel even with a 1,000 ms exposure time. To reduce this saturation and nonspecific staining of viable cells, the final concentration of the viability assay reagent was lowered in subsequent experiments.

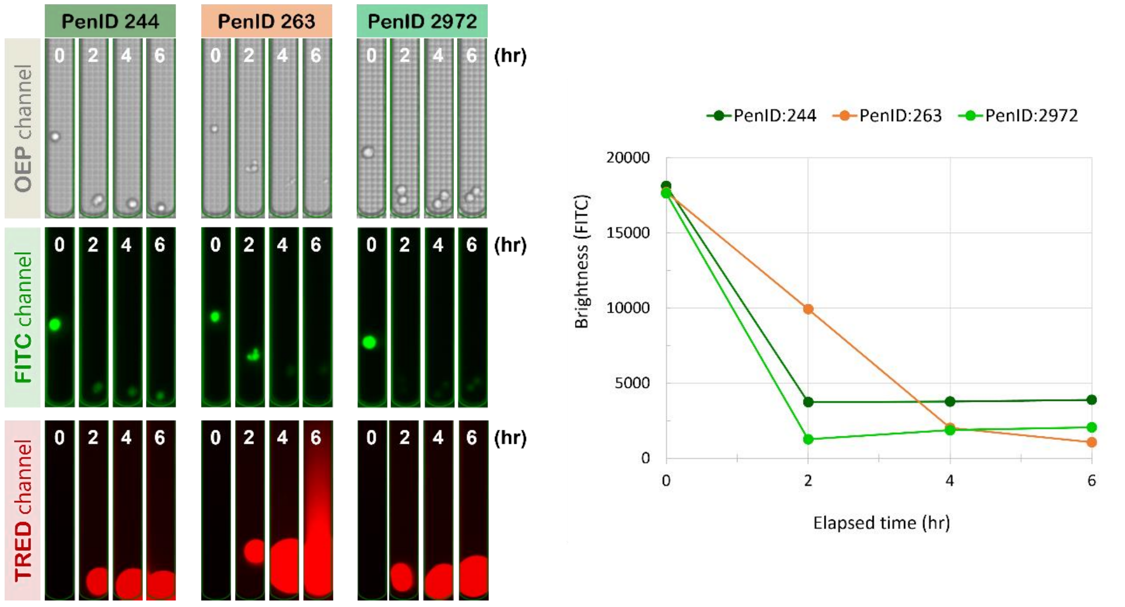

Figure S17. At the 6-hour time point, nonspecific staining by the viability assay reagent was observed in the TRED channel; however, live and dead cells could be clearly distinguished based on morphological differences in bright-field imaging using the OEP channel. Cell death was observed in less than 1% of cells (e.g., PenID 263), characterized by the loss of cellular boundaries in the OEP channel and widespread fluorescence from the viability assay reagent throughout the nanopen in the TRED channel. During the observation period, differences in cellular behavior were noted at the single-cell level, including cells that underwent division and those that did not, as well as cells exhibiting either rapid or gradual decreases in EGFP fluorescence intensity. By the end of the 6-hour culture, no cells exhibited detectable fluorescence in the FITC channel.

### 7. 1:1 co-culture of OBOC-DEL beads and cells

#### 7-1. Workflow for phenotypic screening on Beacon

Based on the previous investigations, OBOC-DEL beads bound to the target compound were individually loaded into nanopens arranged on a microfluidic chip. Subsequently, single cells to be evaluated were loaded into nanopens containing the beads. Upon UV (390 nm) irradiation, the linker was cleaved from the beads, followed by time-lapse imaging over a defined period to monitor phenotypic changes in the cells, particularly changes in EGFP fluorescence intensity. The images obtained from time-lapse imaging were analyzed using image analysis software, and data regarding cell morphology and EGFP fluorescence intensity were collected. Based on the analysis results, particles from nanopens containing target beads were retrieved and unloaded into a 96-well plate. The barcode regions of the recovered beads were amplified by PCR and sequenced using next-generation sequencing (NGS) (Figure 18). Furthermore, the workflow for phenotype-based screening using a 1:1 co-culture format of OBOC-DEL beads and single cells was optimized as shown in Table S3.

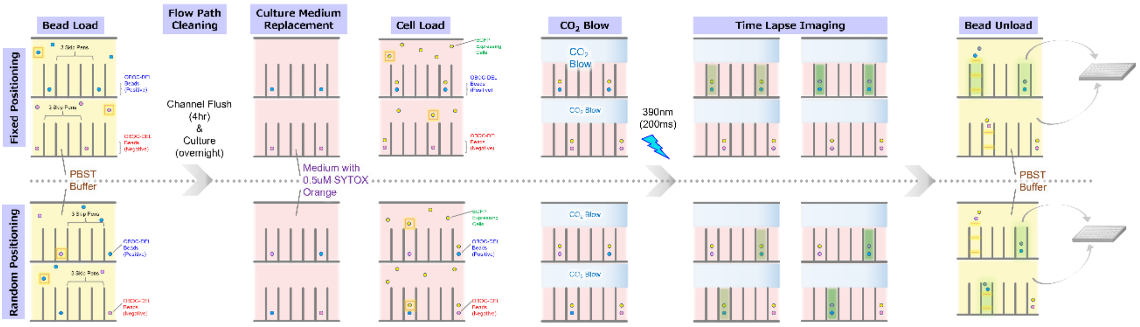

Figure S18. Procedure for Phenotypic Screening via 1:1 Co-culture of OBOC-DEL Beads and Cells Using Beacon. A single cell was loaded into a nanopen containing one OBOC-DEL bead. To prevent compound diffusion from the nanopen, the medium in the channel was removed. After UV (390 nm) irradiation, cells were observed for 6 hours, and nanopens were selected based on the presence or absence of changes in cell brightness. Beads from the selected nanopens were then unloaded and subjected to NGS analysis.

Table S3. Optimized Beacon Workflow and Settings for 1:1 Co-culture of OBOC-DEL Beads and Cells for Phenotypic Screening

| Process | Operations | Details |
|---|---|---|
| Preparation | Prime Line and Chip w/Media | The fluid inside the chip containing the channels and nanopens was replaced with PBST solution which was used to condition the beads. |
| Beads Load | TPS Load | Concentration: 1.0 x $10^6$ beads/mL (set lower than 1.5 x $10^6$ to enable pen skipping)<br>Import beads to chip and pen a single bead to nanopen.<br>Pen Chip x 3 (with 3-skip pens) - Unpen Multiple |
| Channel Cleaning | - Extended Flush<br>- Culture | - 4 hours: Extended Flush ×24 cycles<br>- Culture with PBST buffer (overnight at 25 °C) |
| Replacement with Culture Medium | Prime Line and Chip w/Media | The solution exchange channel and chip were replaced with culture medium (from PBST to cell culture medium supplemented with SYTOX Orange). |
| Cells Load | - TPS Load<br>- Flush | - Single cell penned into nanopens already containing one bead (Pen Chip x 2, 1:1 bead-to-cell ratio per pen)<br>- Concentration: 1.5 x $10^6$ cells/mL (>98% viability, filtered with 40 μm strainer just before the loading step)<br>- Post-load wash: Flush x 3 |
| $CO_2$ Blow | Manual operation | Inject $CO_2$ into the channel of the chip and remove as much liquid as possible from the channel (>30 min).<br>- Blow stopped immediately before assay. |
| Linker photoinduced dissociation & Time Lapse Imaging | - Multi-Spectral Time Lapse Imaging<br>- TPS Imaging | UV (390 nm) irradiation applied to each FOV on the chip; observe FITC fluorescence attenuation in EGFP-expressing cells due to released compounds.<br>Workflow: Chip calibration - TPS Imaging (OEP, FITC: 2,000 ms; TRED: 1,000 ms) - UV irradiation (200 ms) - Time Lapse imaging every 2 hours for 6 hours (OEP, FITC: 2,000 ms; TRED: 1,000 ms) |
| Cleaning | - Flush<br>- Culture | Flow PBST through the channel and chip again. |
| Analysis | - | - Analysis using Image Analyzer (cell death exclusion - positive bead screening).<br>- Image data (e.g., brightness) organized in Excel; hits verified by comparing with images. |
| Beads Unload | Unload | Selected beads were unpenned from the nanopen and transferred to the 96-well collection plate. |

### 7-2. Criteria for selecting target beads

To objectively distinguish between positive and negative classifications in cellular phenotype evaluation, criteria for judgment were established. The evaluation was conducted in nanopens containing one bead and one cell. Prior to the assay, cells were confirmed to be viable and to exhibit clear EGFP expression, as verified using the viability assay reagent in the TRED channel and morphological observation in the brightfield OEP channel. Following a single 200 ms UV irradiation at 390 nm, cells were cultured for 6 hours, and time-dependent phenotypic changes were assessed. Only cells that maintained clear boundaries (OEP channel) and showed no diffusion of fluorescence from the viability assay reagent into the nanopen (TRED channel) after the assay were included in the analysis. Under these experimental conditions, cells exhibiting a decrease in EGFP fluorescence intensity of more than 5,000 in the FITC channel between 0 and 6 hours were classified as positive (Figure 19).

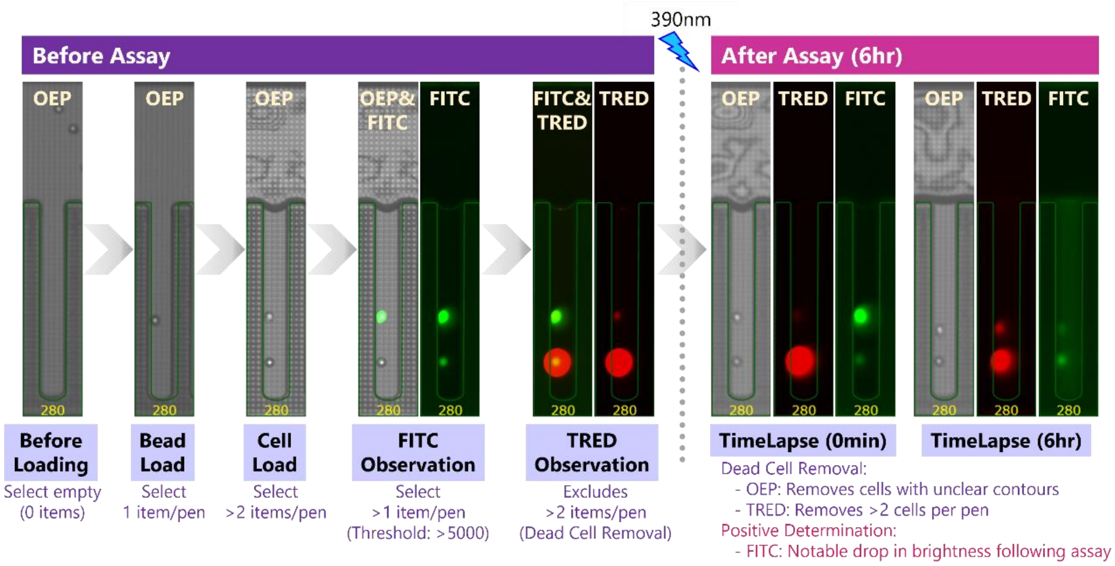

Figure S19. Criteria for Screening and Positive Hit Identification. Nanopens containing a single OBOC-DEL bead and one cell were evaluated based on EGFP fluorescence intensity reduction. Pens showing a significant decrease in EGFP intensity after the assay were classified as positive, whereas those with unchanged fluorescence were classified as negative. Beads from these nanopens were then unloaded and subjected to barcode analysis.

### 7-3. Phenotypic screening of 1:1 OBOC-DEL bead–cell co-cultures

Each nanopen was loaded with a single bead and a single cell. After irradiation with near-UV light at 390 nm through a DAPI filter, the cells were co-cultured for six hours. Imaging was performed every two hours throughout the assay. At the end of the experiment, more than 99% of the cells maintained clear boundaries, and no diffusion of fluorescence derived from the viability dye was observed within the nanopens. These findings confirmed that the cells remained stably viable throughout the assay period. Observation using the FITC channel revealed two distinct cellular phenotypes. Some cells exhibited a reduction in EGFP fluorescence intensity, whereas others showed little to no change in brightness even after six hours (Figure 20). To investigate whether these phenotypic differences were associated with the type of compound present in each nanopen, beads were retrieved from nanopens exhibiting either phenotype. These beads were then subjected to PCR amplification and sequencing analysis to identify the compounds bound to each bead.

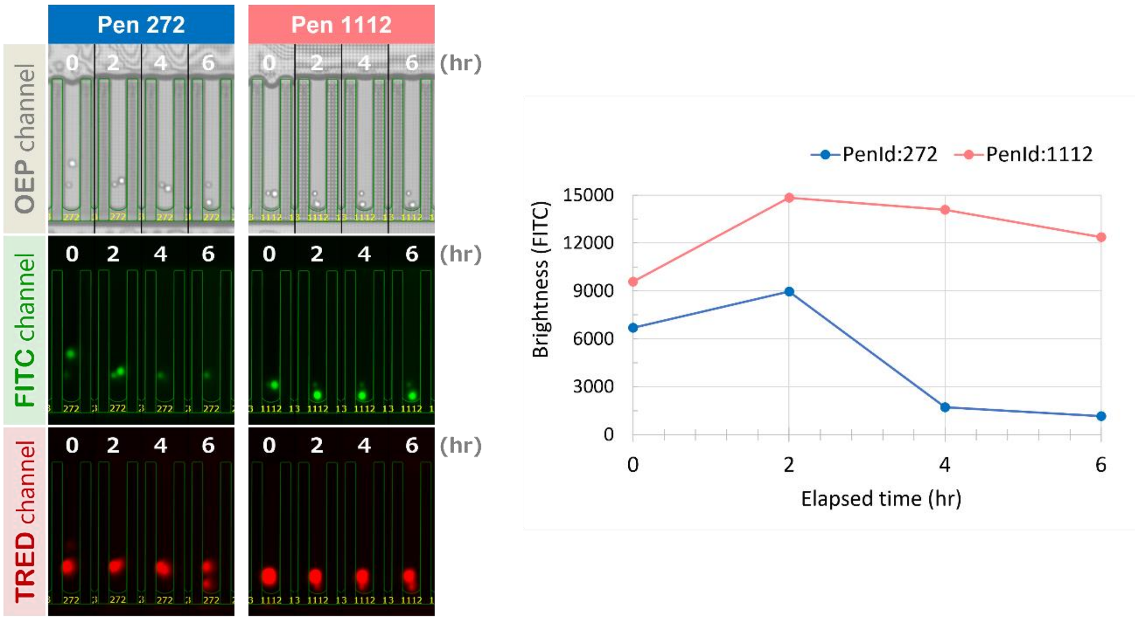

Figure S20. Left: Phenotypic screening of 1:1 OBOC-DEL bead–cell co-cultures using Beacon. Each nanopen containing one bead and one cell was imaged every two hours following UV irradiation. More than 99% of the cells exhibited clear boundaries in the OEP channel, and no diffusion of fluorescence derived from the viability assay reagent was observed in the TRED channel. These findings suggest that the cells remained stably viable within the nanopen. In FITC channel observations, some cells showed a decrease in EGFP fluorescence intensity, while others exhibited little to no change even after 6 hours. Right: Time-dependent changes in EGFP fluorescence intensity. After 6 hours of culture, PenID 272 showed a marked decrease in brightness, whereas PenID 1112 exhibited no significant change.

### 7-4. Recovery of control beads for NGS analysis

As a control for NGS analysis, OBOC-DEL beads were loaded and unloaded without co-culture with cells (Figure S21). Beads used for analysis included individually recovered positive beads (compound-bound), negative beads (unbound), and a mixture of both types collected into a single well at a 1:1 ratio.

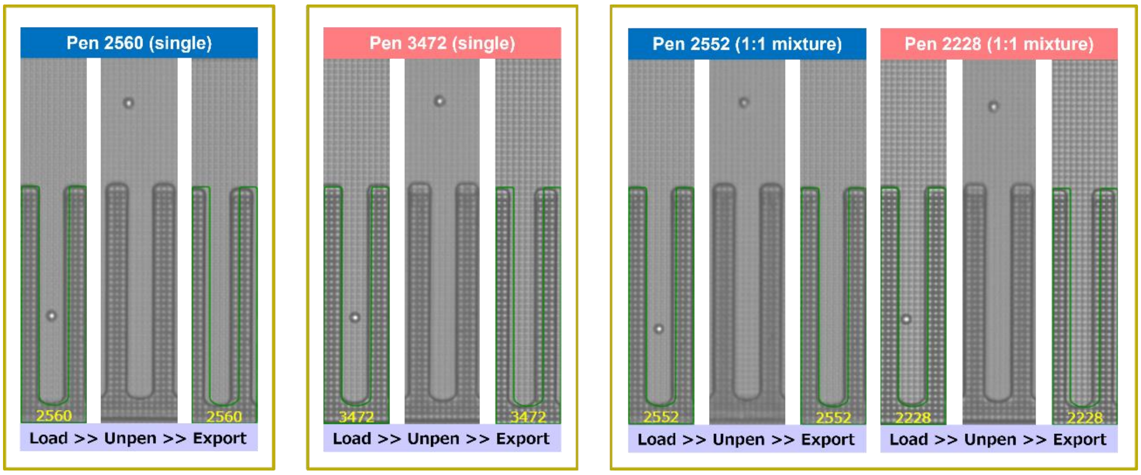

Figure S21. Recovery of Control Beads for NGS Analysis. As a control for NGS analysis, selected OBOC-DEL beads were loaded into designated nanopens without performing co-culture assays with cells. Specifically, Pen #2560 was collected as a positive single bead (compound-bound), and Pen #3472 as a negative single bead (compound-unbound). Furthermore, a positive bead (Pen #2552) and a negative bead (Pen #2228) were simultaneously unloaded, mixed at a 1:1 ratio, and collected into the same well. These beads were subsequently subjected to PCR amplification of the target barcode region and included in the NGS analysis along with the beads screened during the assay.

### 7-5. PCR amplification and sequencing

The each recovered bead (Pen #272 and Pen #1112) and each control bead, including a positive single bead (Pen #2560), negative single beads (Pen #3472, Pen #3472), a mixture of a positive bead (Pen #2552), and a negative bead (Pen #2228), was amplified by two rounds of PCR as previously reported[4]. The control beads (Library control, Positive control, and Negative control), consisting of 1,000 beads, were subjected to two rounds of PCR amplification. The resulting PCR products were sequenced using Illumina high-throughput sequencing, and the obtained data were processed and analyzed according to established protocols[5].

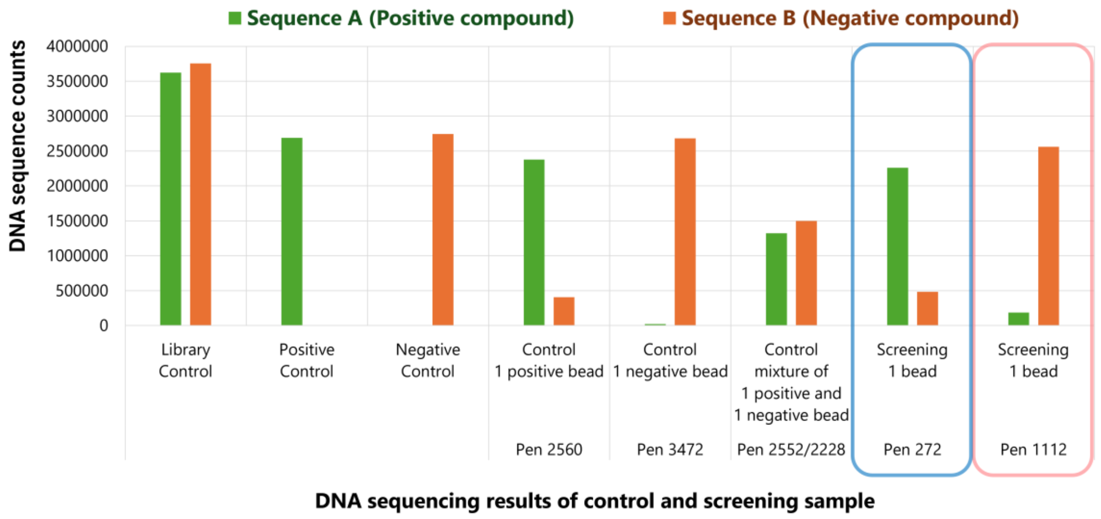

Figure S22. Graph of DNA sequencing results: a) Library control, b) positive control, c) negative control, d) C5 2560, e) D5 3472, f) E5 2552/2228, g) C5 272, h) D5 1112

### 8. References

[1] a) Hockemeyer, D., Soldner, F., Beard, C., Gao, Q., Mitalipova, M., DeKelver, R.C., Katibah, G.E., Amora, R., Boydston, E.A., Zeitler, B., Meng, X., Miller, J.C., Zhang, L., Rebar, E.J., Gregory, P.D., Urnov, F.D., Jaenisch, R. Efficient targeting of expressed and silent genes in human ESCs and iPSCs using zinc-finger nucleases. *Nat. Biotechnol.* **27**, 851–857 (2009). https://doi.org/10.1038/nbt.1562

b) Zotova, A., Lopatukhina, E., Filatov, A., Khaitov, M. & Mazurov, D. Gene editing in human lymphoid cells: role for donor DNA, type of genomic nuclease and cell selection method. *Viruses* 9, 325 (2017). https://doi.org/10.3390/v9110325

[2] a) MacConnell, A. B., McEnaney, P. J., Cavett, V. J., Paegel, B. M. DNA-encoded solid-phase synthesis: encoding language design and complex oligomer library synthesis. *ACS Comb. Sci.* **17**, 518–534 (2015). https://doi.org/10.1021/acscombsci.5b00106

b) Dixit, A., Paegel, B. M. Solid-phase DNA-encoded library synthesis: a master builder's instructions. *Nat. Protoc.* **19**, 1–40 (2025). https://doi.org/10.1038/s41596-025-01190-4

[3] a) Cochrane, W. G., Malone, M. L., Dang, V. Q., Cavett, V., Satz, A. L., Paegel, B. M. Activity-based DNA-encoded library screening. ACS Comb. Sci. 21, 425–435 (2019). https://doi.org/10.1021/acscombsci.9b00037

[4] a) Decurtins, W., Wichert, M., Franzini, R.M., Buller, F., Stravs, M.A., Zhang, Y., Neri, D., Scheuermann, J. Automated screening for small organic ligands using DNA-encoded chemical libraries. *Nat. Protoc.* **11**, 764–780 (2016). https://doi.org/10.1038/nprot.2016.039

[5] a) Wichert, M., Krall, N., Decurtins, W., Franzini, R.M., Pretto, F., Schneider, P., Neri, D. & Scheuermann, J. Dual-display of small molecules enables the discovery of ligand pairs and facilitates affinity maturation. *Nat. Chem.* **7**, 241–249 (2015). https://doi.org/10.1038/nchem.2158
